# Loss of O-GlcNAcylation on MeCP2 Thr 203 Leads to Neurodevelopmental Disorders

**DOI:** 10.1101/2020.06.17.158311

**Authors:** Juanxian Cheng, Zhe Zhao, Liping Chen, Ruijing Du, Yan Wu, Qian Zhu, Ming Fan, Xiaotao Duan, Haitao Wu

## Abstract

Mutations of the X-linked methyl-CpG-binding protein 2 (*MECP2*) gene in humans are responsible for most cases of Rett syndrome (RTT), an X-linked progressive neurological disorder. While genome-wide screens in clinical trials reveal several putative RTT-associated mutations on *MECP2*, their causative relevance regarding the functional regulation of MeCP2 on the etiologic sites at the protein level require more evidence. In this study, we demonstrate that MeCP2 is dynamically modified by O-linked-β-N-acetylglucosamine (O-GlcNAc) at threonine 203 (T203), an etiologic site in RTT patients. Disruption of the O-GlcNAcylation of MeCP2 specifically at T203 impairs dendrite development and spine maturation in cultured hippocampal neurons, and disrupts neuronal migration, dendritic spine morphogenesis and dysfunction of synaptic transmission in the developing and juvenile mouse cerebral cortex. Mechanistically, genetic disruption of O-GlcNAcylation at T203 on MeCP2 decreases neuronal activity-induced induction of *Bdnf* transcription. Our study highlights the critical role of MeCP2 T203 O-GlcNAcylation in neural development and synaptic transmission potentially *via* BDNF.

## INTRODUCTION

Rett syndrome (RTT) is a severe X-linked neurodevelopmental disorder that preferentially occurred in females, with an approximate incidence of 1:10,000 ^1, 2^. Almost 90% of RTT cases are caused by methyl-CpG-binding protein 2 (*MECP2*) gene mutations such as missense, nonsense, insertion, deletion, and splice-site variations ^3, 4, 5^, and loss of MeCP2 is closely related to the occurrence of RTT ^1, 6, 7, 8^. MeCP2 activity in normal central nervous system (CNS) development and function is controlled by both precise expression levels ^9, 10^ and post-translational modifications (PTMs). For example, neurons from *Mecp2*-null mutant mice have smaller somas ^11, 12^, decreased dendritic complexity ^13, 14, 15^ and dysfunction of synaptic plasticity ^16, 17, 18^. Gain-of-function MeCP2 by overexpression in transgenic mice or monkeys results in progressive neurological and psychiatric dysfunctions ^19, 20^. These genetic studies suggest that precise and dynamic expression of MeCP2 is critical to maintain normal brain development and function.

In addition to expression level, PTMs on MeCP2 such as phosphorylation have been demonstrated as critical regulators of its role in dendritic growth, spine maturation, and activation of calcium-dependent brain-derived neurotrophic factor (BDNF) gene expression, suggesting that MeCP2 PTMs particularly impact neurodevelopment processes and activity-dependent gene expression. ^21, 22, 23, 24, 25^. Recently, the novel PTM O-GlcNAcylation has emerged as a potent regulator of neurogenesis and synaptic plasticity ^26, 27, 28^. O-GlcNAcylation is a highly dynamic process ^29, 30, 31^. O-GlcNAc transferase (OGT) catalyzes the addition of O-GlcNAc to serine and threonine residues on intracellular proteins, whereas O-GlcNAcase (OGA) results in the removal of O-GlcNAc modifications ^29, 31, 32^. Previous studies have identified O-GlcNAcylation as a potent modulator of neuronal differentiation ^26, 33^ and synaptic plasticity ^34^. Interestingly, MeCP2 was found to be O-GlcNAcylated and phosphorylated simultaneously in rat cortical neurons ^35, 36^. However, despite the critical role of both O-GlcNAcylation and MeCP2 on neural development and synaptic transmission, the physiological function and molecular mechanisms of human MeCP2 O-GlcNAcylation remain elusive.

In this report, we first used mass spectrometry (MS) to systematically identify O-GlcNAcylation sites on mice, rats, and human MeCP2. We found that human MeCP2 T203, a site previously implicated in the pathogenesis of RTT ^37, 38, 39^, was O-GlcNAcylated at relatively high levels at baseline compared to rodent species. Furthermore, we demonstrated the critical role of MeCP2 T203 O-GlcNAcylation in the regulation of dendrite outgrowth, dendritic spine morphogenesis and synaptic transmission both *in vitro* and *in vivo*. Mechanistic studies suggest that this may be due to the regulation of neuronal activity-induced induction of *Bdnf* transcription. Together, our results identified a previously unknown function of O-GlcNAcylated MeCP2 T203, which may be essential for understanding the molecular mechanisms behind the neuropathology of *MECP2* mutation-caused RTT.

## RESULTS

### Identification of O-GlcNAcylation Sites on MeCP2

To understand whether MeCP2 is a highly conserved protein in different species, we first investigated the evolutionary divergence of the *Mecp2* gene from humans and 25 additional representative species using phylogeny tree reconstruction. We found that the protein sequences of MeCP2 are conserved among most mammals, with distance values less than 0.1, suggesting the sequence similarity between each pair of mammalian MeCP2 protein is over 90%. In contrast, sequence distances were much higher between mammalian and non-mammalian species. This indicates that the protein sequence and function of MeCP2 have undergone substantial changes during the evolution of vertebrates, but are relatively conserved in mammals (Fig. 1A).

**Figure 1.**
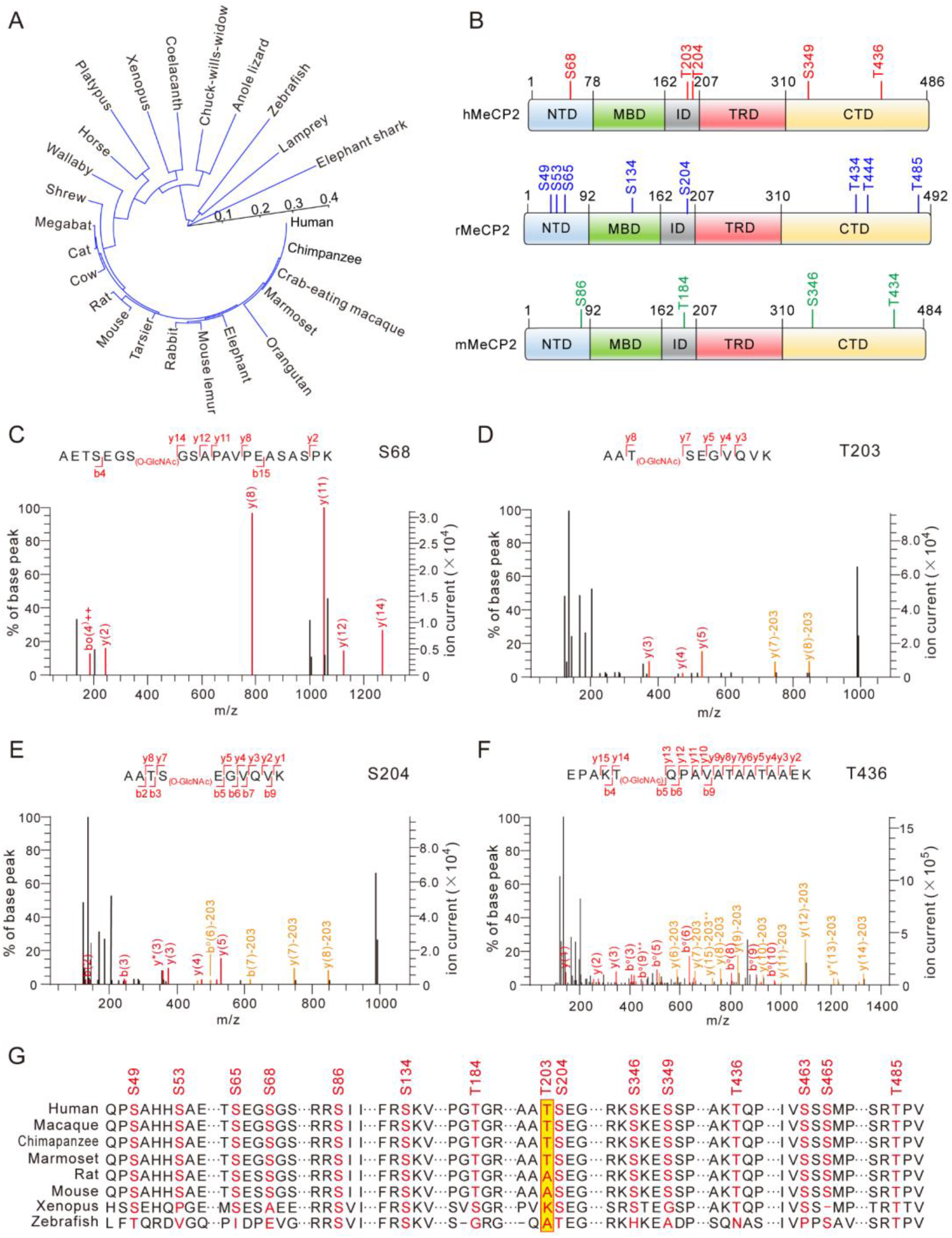
Identification of O-GlcNAcylation Sites in Rodent and Human MeCP2. (A) The phylogeny tree of MeCP2 protein sequence from 25 representative species. The length of each branch represents the evolutionary distance between the MeCP2 protein sequences. The dashed scale line indicates the values of evolutionary distance. (B) MS identified O-GlcNAcylation sites in rodent and human MeCP2 protein. hMeCP2, human MeCP2; rMeCP2, rat MeCP2; mMeCP2, mice MeCP2; NTD, the N-terminal domain; MBD, the methyl-CpG binding domain; ID, the intervening domain; TRD, the transcriptional repression domain; CTD, the C-terminal domain. (C-F) Four representative mass spectra of O-GlcNAcylation sites in hMeCP2 are shown. hMeCP2 was purified from HEK293T cells and analyzed by MS to identify the O-GlcNAcylation sites. (G) Multiple-sequence alignment of MeCP2 protein to reveal the conservation of O-GlcNAcylation site. The O-GlcNAcylation sites in MeCP2 from eight different representative species are shown. The identified O-GlcNAcylation sites are shown in red, and T203 site is highlighted in yellow.

O-GlcNAcylation is highly dynamic and reversible in cellular systems. In order to better capture and concentrate O-GlcNAcylated MeCP2 for MS, HA-tagged MeCP2 in different species were co-expressed with OGT in HEK293T cells, and co-immunoprecipitated (co-IP) with anti-HA antibody-conjugated beads followed by in-gel trypsin digestion (Fig. S1A). The digested MeCP2 peptides were subjected to high-resolution mass spectrometry (MS, nanoLC-LTQ-CID/ETD-Orbitrap) analysis (Table S1) (Fig. S1B-F). In addition to the O-GlcNAcylated sites on T434 and T444, which have been previously identified in rat MeCP2 ^35, 36^, we mapped 11 novel O-GlcNAcylation sites from mouse (mMeCP2), rat (rMeCP2) and human (hMeCP2) (Fig. 1B). To examine the potential relevance of these novel MeCP2 O-GlcNAc sites for neurodevelopmental disorders including RTT, we next compared them with the RTT disease-related *MECP2* mutation sites reported from RettBASE ^37, 38, 39^, a database that catalogues clinically-relevant *MECP2* mutations. Interestingly, we found that three O-GlcNAcylated sites (S68, T203 and S204) were mutated in some RTT clinical cases ^40^ (Table S2). In addition, human MeCP2 T436 (homologue of T434 in mMeCP2 and rMeCP2) O-GlcNAcylation may also be implicated, although no identified clinical cases have been associated with the mutation. Therefore, we mainly focused on four O-GlcNAcylated sites of human MeCP2 including S68, T203, S204 and T436 in the following assays (Fig. 1C-F). Of these sites, additional analysis pointed to T203 as a particularly interesting target. We compared the MeCP2 O-GlcNAcylation sites identified in this study from eight different species to assess their conservation (Fig. 1G). While the majority of these sites are conserved among vertebrate animals, T203 O-GlcNAcylation was selectively conserved in primates such as human, macaque, chimpanzee, and marmoset, but not in rodents. Given that MeCP2 T203 is an etiologic site in RTT patients, the T203 site may carry particular evolutionary functional significance in the brains of high primates.

### MeCP2 is O-GlcNAcylated at T203 by Direct Interaction with OGT

The above MS results indicate that MeCP2 may serve as a new protein substrate of OGT (Fig. S1G). To test this hypothesis, we first investigated the interaction between MeCP2 and OGT, and to identify the MeCP2-binding domain within OGT. Endogenous MeCP2 from mouse whole brain lysates were co-immunoprecipitated with native OGT (Fig. 2A). Reciprocal assays in mouse brain lysates showed that OGT also co-immunoprecipitated with MeCP2 (Fig. 2B), strongly suggesting physiological interaction *in vivo*. To further explore if MeCP2 and OGT bind via direct protein-protein interaction, we performed an *in vitro* GST protein pull-down assay using purified GST-fused OGT and human MeCP2 protein. GST-fused OGT, but not GST alone, was able to pull down MeCP2 and *vice versa*, suggesting direct binding with each other (Fig. 2C and D). To reveal the reciprocal binding domains supporting the interaction of OGT with MeCP2, we first generated various human OGT deletion mutants. Co-IP assays showed that deletion of the entire N-terminal TPR domain (aa 2-465) of OGT completely disrupted the interaction between OGT and MeCP2, suggesting the TPR domain is necessary (Fig. 2E and F). Next, domain mapping analysis showed that deletion of 4-6 TPR (aa 114-214), 7-9 TPR (aa 215-316) or 10-13.5 TPR (aa 317-465), had no effects on the interaction between OGT and MeCP2 (Fig. 2E and F). In contrast, deletion of the 1-3 TPR domain (aa 2-113) of OGT significantly abolished the binding (Fig. 2E-H), suggesting OGT directly interacts with MeCP2 *via* its N-terminal 1-3 TPR domain. Moreover, in order to identify critical OGT-binding domains on MeCP2, we used MeCP2 deletion mutants in co-IP binding experiments with OGT, and found that both NTD and CTD domains of MeCP2 are necessary (Fig. S2).

**Figure 2.**
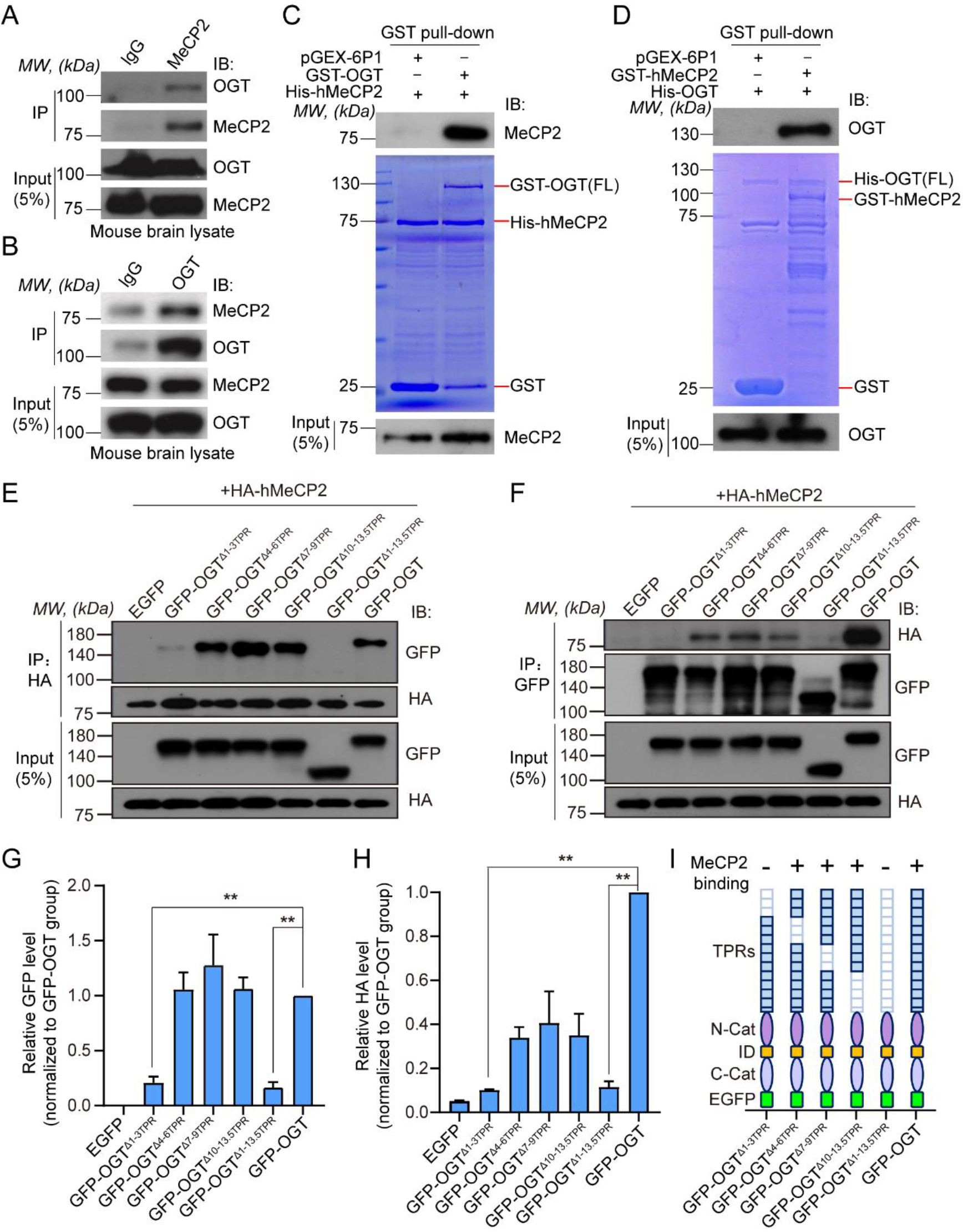
OGT Directly Interacts with and O-GlcNAcylates MeCP2. (A) The mouse brain lysates were immunoprecipitated with an anti-MeCP2 antibody, followed by Western blot analysis with an anti-OGT antibody. (B) Reciprocal co-IP assay of mouse brain lysates with an anti-OGT antibody, followed by Western blot analysis with an anti-MeCP2 antibody. (C-D) GST pull-down assays for His-hMeCP2 and GST-OGT, or His-OGT and GST-hMeCP2. Input and pull-down samples were analyzed with anti-MeCP2 and anti-OGT antibodies. Input represents 5% of the amount used for pull-down assay. (E-F) Cell lysates from HEK293T cells transfected with HA-tagged hMeCP2 and GFP-tagged full length and indicated deletion mutants of OGT were immunoprecipitated with anti-HA or anti-GFP antibodies, followed by Western blot analysis with anti-GFP and anti-HA antibodies, respectively. Input represents 5% of the amount used for pull-down assay. (G-H) Representative quantification of Western blot results followed by co-IP assay in E and F, respectively. Histograms show mean ± SEM. One-way ANOVA followed by Bonferroni test, **p < 0.01. (I) A summarized diagram for the serial deletion mutants of OGT and their binding capacities to MeCP2 is shown. Deletion of either the entire TPRs or 1-3 TPRs domain within OGT dramatically disrupts its binding to MeCP2.

To further study the mechanisms of MeCP2 O-GlcNAcylation by OGT *in vivo*, we setup a well-established chemoenzymatic labelling approach ^41^ to detect MeCP2 O-GlcNAcylation from whole brain lysates (Fig. 3A). O-GlcNAc-modified proteins from mouse brain lysates were enzymatically labelled with an azido-N-acetylgalactosamine sugar and biotinylated via Cu(I)-mediated [3+2] azide-alkyne cycloaddition (CuACC) chemistry, which were captured with streptavidin agarose beads, and subsequently immunoblotted with an antibody against MeCP2 (Fig. 3A). As shown in Fig. 3B, O-GlcNAcylation occurs in endogenous MeCP2 in wild type (WT) mice *in vivo*, and much stronger O-GlcNAcylation modification of MeCP2 was detected in overexpression of *MECP2* transgenic mice ^19^ compared with WT mice. The above results suggest direct protein-protein binding between MeCP2 and OGT that results in O-GlcNAcylation of MeCP2 (Fig. 2). To further confirm that OGT catalyzes MeCP2 upon direct protein binding, we used co-IP assay to demonstrate that ectopic GFP-tagged hMeCP2 was O-GlcNAcylated in the presence of OGT in transfected HEK293T cells (Fig. 3C). Additionally, *in vitro* O-GlcNAcylation assay further confirmed that His-tagged hMeCP2 was directly O-GlcNAcylated by GST-tagged enzyme domain of OGT (323-1041) ^42^ (Fig. 3D).

**Figure 3.**
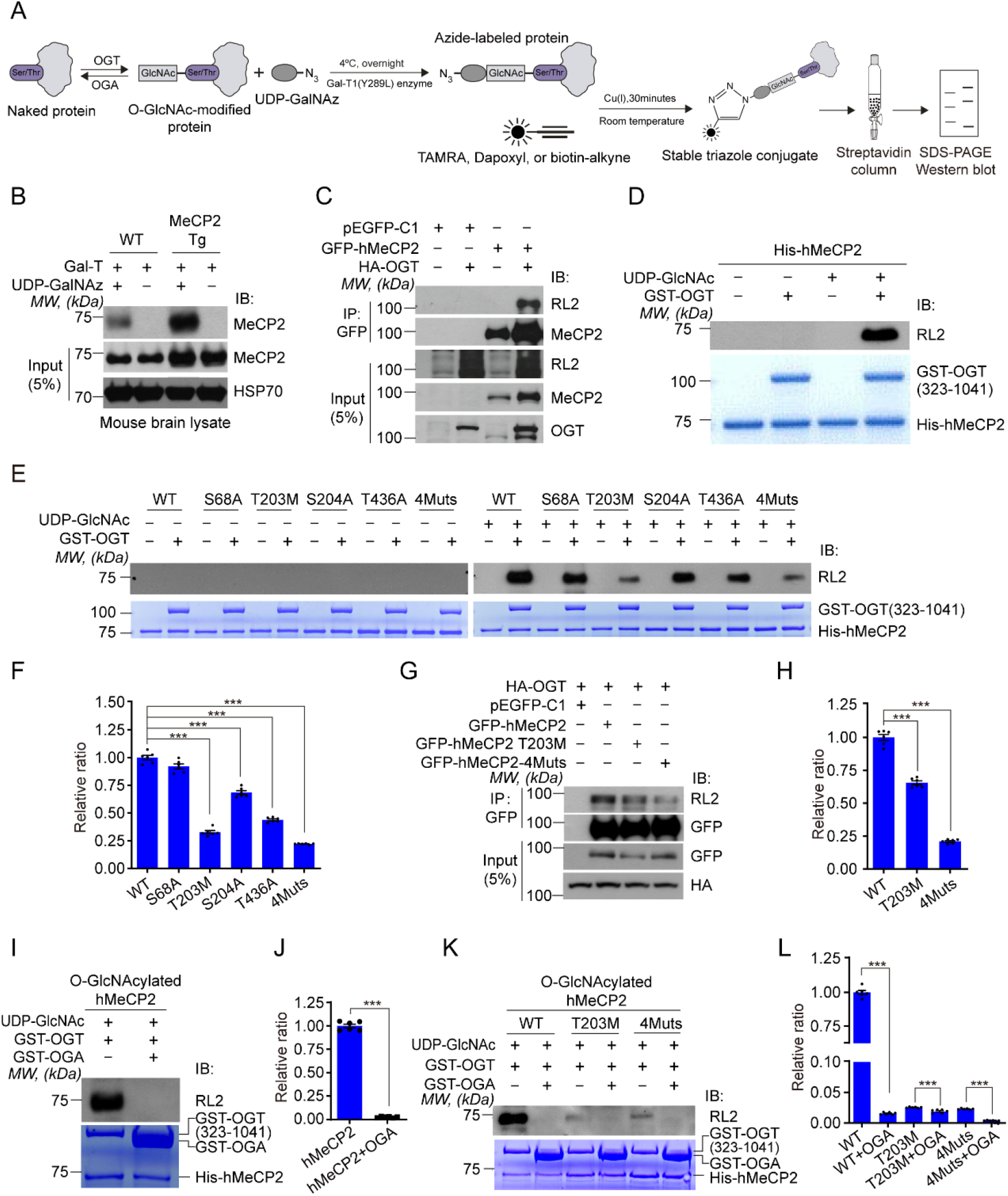
T203 Residue of hMeCP2 is Dynamically O-GlcNAcylated by OGT. (A) schematic depicting the chemoenzymatic labelling approach for biotinylation, capture, and detection of O-GlcNAcylated protein from brain or cell lysates. (B) Detection of O-GlcNAcylated MeCP2 protein in brain lysates from wild type (WT) and *MECP2* transgenic (Tg) mice using chemoenzymatic labelling approach. Higher level of O-GlcNAcylated MeCP2 was detected in *MECP2* Tg mice compared with WT control. In the absence of GalT or UDP-GalNAz, no O-GlcNAcylated MeCP2 can be detected. HSP70 was used as loading control. (C) OGT elevates the O-GlcNAcylation level of MeCP2. Exogenous GFP-tagged hMeCP2 was co-expressed with or without HA-tagged OGT in HEK293T cells for co-IP assay followed by Western blot analysis with anti-MeCP2 and anti-RL2 antibodies, respectively. Input represents 5% of the amount used for co-IP assay. pEGFP-C1 mock vector was used as a negative control. (D) OGT directly O-GlcNAcylates MeCP2 by *in vitro* glycosylation assay. Recombinant His-tagged hMeCP2 protein was incubated with or without purified GST-OGT (323-1041) and UDP-GlcNAc, followed by Western blot analysis with an anti-RL2 antibody. The loading amount of recombinant proteins used for the *in vitro* assay was confirmed by Coomassie blue staining. (E-F) Identification of the O-GlcNAc modified sites in MeCP2 by *in vitro* glycosylation assay. Purified wild type or mutant His-hMeCP2 was used as substrates of GST-OGT (323-1041) in the presence of UDP-GlcNAc. The loading amount of GST-OGT and His-hMeCP2 was confirmed by Coomassie blue staining. Quantification analysis shows the significantly decreased O-GlcNAc level in the hMeCP2 mutations including T203M, S204A, T436A and 4Muts. Histograms show mean ± SEM. One-way ANOVA followed by Bonferroni test, ***p < 0.001. (G-H) Co-IP assay shows dramatically reduced the O-GlcNAc modified level in T203M or 4Muts of hMeCP2 compared with WT in the presence of OGT in transfected HEK293T cells. Cell lysates were immunoprecipitated with anti-GFP beads and immunoblotted as indicated in G. Input represents 5% of the total amount used for co-IP assay. pEGFP-C1 mock vector was used as a negative control. Quantification analysis shows the significantly decreased RL2 level in T203M and 4Muts mutants compared with wild type control. Histograms show mean ± SEM. One-way ANOVA followed by Bonferroni test, ***p < 0.001. (I-J) OGA reversely regulates the O-GlcNAcylation of hMeCP2 by *in vitro* glycosylation assay. Quantification analysis shows the significantly decreased RL2 level of hMeCP2 in the presence of GST-OGA. Histograms show mean ± SEM. *t*-test, ***p < 0.001. (K-L) The O-GlcNAcylation of T203M mutant or 4Muts mutant of hMeCP2 was also reversed by OGA by *in vitro* glycosylation assay. Quantification analysis shows the significantly decreased RL2 level of WT and hMeCP2 mutants in the presence of GST-OGA. Histograms show mean ± SEM. One-way ANOVA followed by Bonferroni test, ***p < 0.001.

Because our MS results indicated several O-GlcNAcylation sites in hMeCP2 with potential relevance to RTT (Fig. 1 and Fig. S1) we wanted to further identify major functional O-GlcNAcylation activity among these sites. To test this, MeCP2 S68A, T203M, S204A, T436A, and quadruple site mutation to alanine (4 Muts) were generated. We found that the level of O-GlcNAcylation was significantly reduced in the T203M and 4 Muts mutant variations *in vitro* (Fig. 3E and F) and *in vivo* (Fig. 3G and H). However, changes in O-GlcNAcylation were insignificant for other MeCP2 mutants, suggesting that the T203 site is the predominant O-GlcNAcylation site on human MeCP2. Of note, the T203 site has previously been implicated in the pathogenesis of RTT. Interestingly, MeCP2-4Muts still showed a weak level of O-GlcNAcylation, which further supports our MS results showing multiple O-GlcNAcylation sites on MeCP2 (Fig. 1 and Fig. S1).

To interrogate the reversibility of MeCP2 O-GlcNAcylation, we introduced GST-tagged OGA (31-624) recombinant protein, which removes O-GlcNAc modifications. OGA almost completely removed the O-GlcNAcylation on MeCP2 *in vitro* (Fig. 3I-L), implying that MeCP2 O-GlcNAcylation is a reversible and dynamic process.

### T203 O-GlcNAcylation is Required for Dendrite Development in Cultured Neurons

To understand the functional importance of MeCP2 O-GlcNAcylation, we next examined its impact on neural development. Overexpression of MeCP2 inhibits dendritic growth of hippocampal neurons ^43, 44^, and *MECP2* transgenic mice also appears to have a progressive neurological and neurobehavioral disorders ^19, 44, 45^. If MeCP2 O-GlcNAcylation is involved in these aberrant neural developmental physiology, we reasoned that mutating O-GlcNAc sites in an MeCP2 overexpression system may rescue the above observed defects. Thus, for our next experiments, we employed MeCP2 overexpression as an experimental model to test the significance of MeCP2 T203 O-GlcNAcylation for neural development.

First, we compared the morphological differences of dendritic branches of cultured mouse hippocampal neurons using anti-MAP2 immunofluorescent staining after infection with the indicated LV-GFP control or LV-hMeCP2 ectopic expression lentivirus (Fig. S3A). The total length of dendrites was significantly inhibited by overexpressing ectopic hMeCP2 compared with cells infected with the LV-GFP control virus (207.4 ± 7.85 μm in LV-hMeCP2 and 294.2 ± 10.17 μm in LV-GFP control, p < 0.001, n = 62 and 67 neurons, respectively). However, overexpression of hMeCP2 T203M or hMeCP2-4Muts mutant had no significant effect on the length of total dendrite compared with LV-GFP control neurons (269.0 ± 9.58 μm in LV-hMeCP2 T203M and 263.6 ± 8.81 μm in LV-hMeCP2-4Muts, respectively; p = 0.1348 and 0.0829, n = 63 and 56 neurons, respectively) (Fig. S3B and C). In stark contrast, overexpression of the hMeCP2 O-GlcNAc mutant hMeCP2 S68A, S204A or T436A all had similar inhibitory effects on dendrite development as WT hMeCP2 (p < 0.001, n =70, 60 and 63 neurons, respectively). These results suggest that T203M, but not S68A, S204A or T436A, specifically rescues the aberrant dendritic morphology seen with hMeCP2 overexpression. In addition, we also quantified the length of primary and secondary dendritic branches after overexpression of the indicated ectopic hMeCP2. As shown in Fig. S3D and E, overexpression of WT hMeCP2 significantly inhibited the length of both primary and secondary branches compared with LV-GFP controls (162.0 ± 6.02 μm in LV-hMeCP2 and 214.4 ± 6.56 μm in LV-GFP control for primary branches, p < 0.001, n = 62 neurons; 31.3 ± 3.70 μm in LV-hMeCP2 and 71.4 ± 5.87 μm in LV-GFP control for secondary branches, p < 0.001, n = 27 neurons). Overexpression of hMeCP2 S68A, S204A and T436A also significantly decreased the length of primary branches (p < 0.001, n = 58, 50 and 63 neurons, respectively), but overexpression of MeCP2 T203M or MeCP2-4Muts mutant had no obvious effect on the length of both primary and secondary branches (Fig. S3D and E). The rescue effect seen with MeCP2 T203M and MeCP2-4Muts seen to be relatively specific, because overexpression of either S68A, S204A or T436A mutant inhibited neuronal dendrite development similar to overexpressed WT hMeCP2 (Fig. S3B-E). Together, these results indicate that O-GlcNAcylation of hMeCP2 T203 critically underlies the dendritic deficits observed with overexpression of ectopic hMeCP2 in cultured hippocampal neurons, and mutating T203 rescues these deficits in dendritic length and branching.

To further illustrate the significance of MeCP2 O-GlcNAcylation in neural development, we tested whether T203 O-GlcNAcylation had any effects on dendritic spine morphogenesis in cultured hippocampal neurons. A lentiviral-based rescue (LEMPRA) construct was used to exogenously introduce Flag-tagged shRNA-resistant human *MECP2* to replace endogenous mouse *Mecp2*, the expression of which was specifically knocked down by shRNA ^24^ (Fig. 4A). Cultured hippocampal neurons from E17.5 mouse embryos were infected with the indicated lentivirus at DIV 7, then fixed at DIV 14 for immunofluorescent staining with the postsynaptic marker PSD-95 antibody (Fig. 4B) to evaluate the formation of dendritic spines (Fig. 4C). Efficient knockdown of endogenous mMeCP2 and expression of equivalent levels of exogenous hMeCP2 were confirmed by Western blot analysis (Fig. S4). Consistent with previous reports showing smaller somas in *Mecp2*-null neurons ^11, 12^, quantification revealed that the area of soma in shMeCP2 (mMeCP2-) neurons is dramatically reduced compared with FUGW-GFP control neurons (129.1 ± 8.18 μm^2^ in mMeCP2-, and 211.5 ± 18.7 μm^2^ in FUGW-GFP control, p < 0.01, n = 22 and 15 neurons, respectively). Importantly, exogenous expression of WT hMeCP2 or hMeCP2 T436A (203.4 ± 16.71 μm^2^ in mMeCP2−/hMeCP2+ and 200.9 ± 16.04 μm^2^ in mMeCP2−/hMeCP2 T436A+, p = 0.9904 and 0.9875, respectively, compared with FUGW-GFP control, n = 20 and 9 neurons, respectively), but not hMeCP2 T203M or hMeCP2-2Muts mutants (162.9 ± 8.43 μm^2^ in mMeCP2−/hMeCP2 T203M+ and 143.9 ± 9.49 μm^2^ in mMeCP2-/hMeCP2-2Muts+, p = 0.0348 and 0.0174, respectively, compared with FUGW-GFP control, n = 23 and 9 neurons, respectively), efficiently rescued the smaller soma deficiency in mMeCP2- neurons (Fig. 4D).

**Figure 4.**
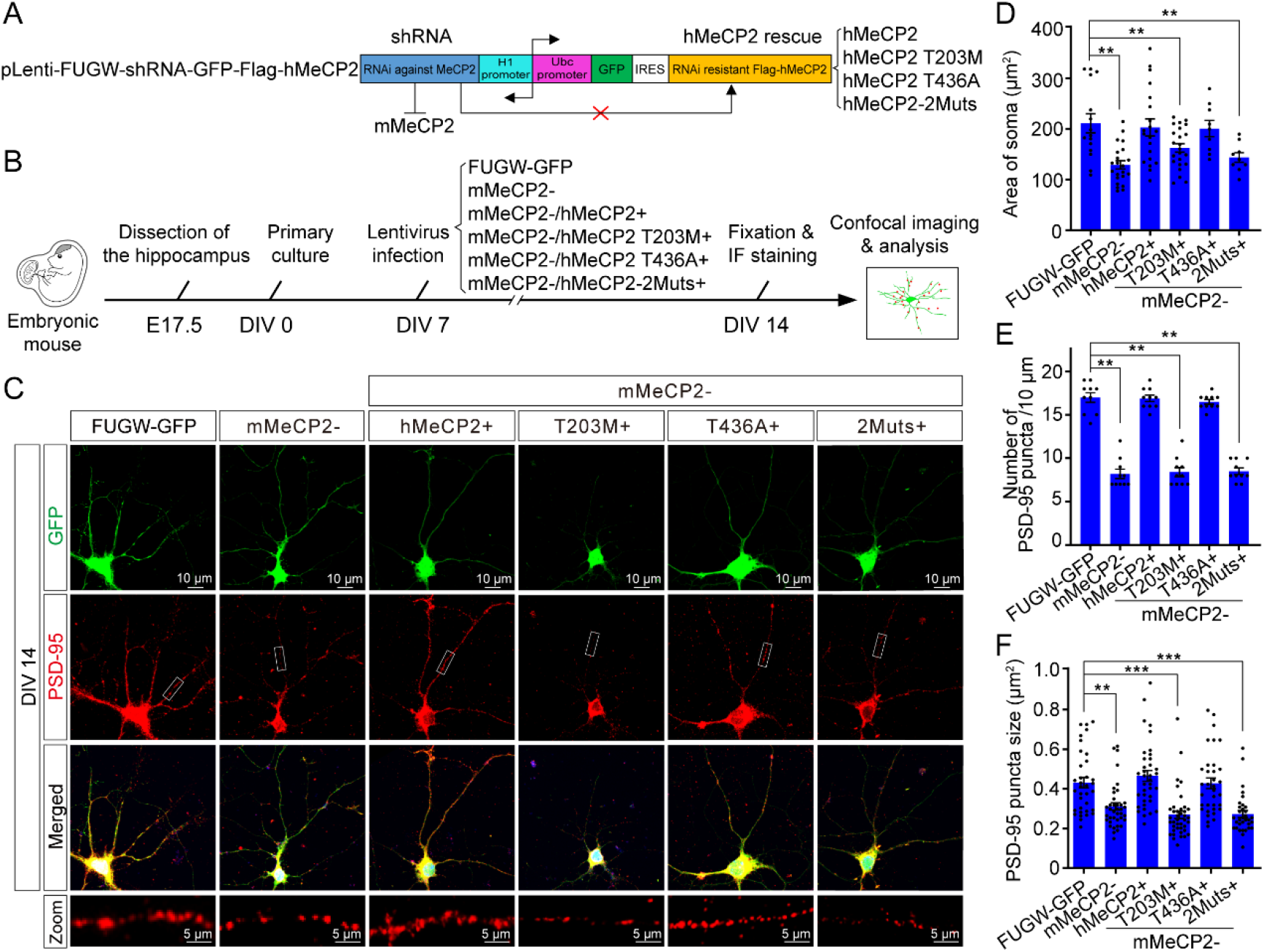
T203 O-GlcNAcylation is Required for Dendritic Spine Formation and Soma Size Maintenance in Cultured Hippocampal Neurons. (A) The recombinant <u>le</u>ntivirus-<u>me</u>diated <u>p</u>rotein-<u>r</u>eplacement <u>a</u>ssay construct, pLEMPRA-MeCP2, was generated to knockdown endogenous mMeCP2 and express Flag-tagged ectopic hMeCP2. (B) Schematic of the experimental design. The primary hippocampal neurons were isolated from E17.5 mouse embryos for *in vitro* culture. The neurons at DIV 7 were infected with the indicated LEMPRA lentivirus for 7 days, and then fixed at DIV 14 for immunofluorescent staining and confocal imaging. (C) Representative pictures of mouse primary hippocampal neurons infected with indicated lentivirus at DIV 7, lentivirus FUGW-GFP was used as negative control. Neurons were collected and fixed at DIV 14 for immunofluorescent staining with anti-PSD-95 antibody for measurement of dendritic spines. The soma and dendrites of indicated LEMPRA lentivirus infected positive neurons were illustrated by GFP (green), and the distribution of PSD-95 positive dendritic spines were shown in red. Boxed areas of PSD-95 positive puncta (red) along the secondary dendritic branches are shown at higher magnification to illustrate the detailed dendritic spine density (Zoom panels). Scale bar represents 10 μm and 5 μm, respectively. (D) Quantification of the area of soma of indicated lentivirus infected positive neurons. Histograms show mean ± SEM. One-way ANOVA followed by Bonferroni test, **p < 0.01. (E-F) Quantification of the linear density and the area of PSD-95 puncta along the secondary dendritic branches in indicated lentivirus infected positive neurons, respectively. Histograms show mean ± SEM. One-way ANOVA followed by Bonferroni test, **p < 0.01, ***p < 0.001.

In agreement with previous reports in *Mecp2*-null mice ^9, 46, 47^, we found that the density of PSD-95 puncta was significantly decreased after knocking down endogenous MeCP2 (8.2 ± 0.53 /10 μm in mMeCP2- and 17.0 ± 0.56 /10 μm in FUGW-GFP control, p < 0.001, n = 10 neurons). Both exogenously expressed Flag-tagged WT hMeCP2 and hMeCP2 T436A mutants efficiently rescued the decrease in PSD-95 puncta after MeCP2 knockdown (16.9 ± 0.39 /10 μm in mMeCP2−/hMeCP2+ and 16.5 ± 0.27 /10 μm in mMeCP2−/hMeCP2 T436A+, p = 0.9997 and 0.8970, respectively, compared with FUGW-GFP control, n = 10). Interestingly, in stark contrast, exogenously expressed hMeCP2 T203M and hMeCP2-2Muts both showed significantly decreased number of PSD-95 puncta compared with FUGW-GFP control neurons (8.4 ± 0.52 /10 μm in mMeCP2−/hMeCP2 T203M+ and 8.5 ± 0.37 /10 μm in mMeCP2-/hMeCP2-2Muts+, p < 0.01 and p < 0.01, respectively, n = 10 neurons) (Fig. 4E), demonstrating an inability to rescue MeCP2 knockdown-induced decrease in PSD-95. In addition, we also analyzed the average size of each PSD-95 puncta within the dendritic spines. We found that the PSD-95 puncta size was significantly decreased in mMeCP2- neurons compared to FUGW-GFP control neurons (0.312 ± 0.018 μm^2^ in mMeCP2-, n = 35; 0.432 ± 0.026 μm^2^ in FUGW-GFP control, n = 38; p = 0.001).

Ectopic expression of WT hMeCP2 and hMeCP2 T436A mutants both significantly rescued the decreased PSD-95-positive puncta size in mMeCP2- neurons to normal level compared with FUGW-GFP control neurons (0.466 ± 0.027 μm^2^ in mMeCP2−/hMeCP2+, n = 37; 0.428 ± 0.027 μm^2^ in mMeCP2−/hMeCP2 T436A+, n = 34; p = 0.738 and 0.999, respectively). However, exogenously expressed hMeCP2 T203M and hMeCP2-2Muts in mMeCP2- neurons both showed significantly decreased size of PSD-95-positive puncta compared with FUGW-GFP control neurons (0.270 ± 0.019 μm^2^ in mMeCP2−/hMeCP2 T203M+, n = 37; 0.273 ± 0.018 μm^2^ in mMeCP2-/hMeCP2-2Muts, n = 33; p < 0.001 and p < 0.001, respectively) (Fig. 4F). Together, these results suggest a requirement of T203 O-GlcNAcylation for dendritic spine morphogenesis. Taken together, our results indicate that hMeCP2 T203 O-GlcNAcylation is essential for the regulation of neurite outgrowth and dendritic spine morphogenesis in cultured hippocampal neurons. Moreover, T203 O-GlcNAcylation is also sufficient for the maintenance of neuronal soma during neuronal differentiation *in vitro*.

### T203 O-GlcNAcylation Promotes Cortical Neuron Migration and Maturation in vivo

To further understand the role of T203 O-GlcNAcylation in cortical development *in vivo*, we manipulated the expression of exogenous human MeCP2 in the developing mouse neocortex using the above-described lentiviral-based rescue (LEMPRA) plasmid by *in utero* electroporation ^24, 48^. Neuronal migration and dendritic differentiation are critical events in cortical construction. Therefore, we analyzed the effects of T203 O-GlcNAcylation on neuronal migration and dendritic spine morphogenesis at embryonic day 17.5 (E17.5) and postnatal day 15 (P15), respectively (Fig. 5A). First, we found that loss-of-function of MeCP2 by shRNA-mediated knockdown resulted in significant migration defects at E17.5 when compared with electroporation of only GFP (Fig. 5B and C). To exclude possible off-target effects of the shRNA system, we showed that the migration defects in MeCP2 knockdown neurons could be rescued by exogenous expression of WT hMeCP2. However, exogenous expression of the hMeCP2 T203M mutant could not rescue the migration defects in mMeCP2 knockdown neurons (Fig. 5B and C), indicating T203 O-GlcNAcylation is required for proper neuronal migration in the embryonic neocortex.

**Figure 5.**
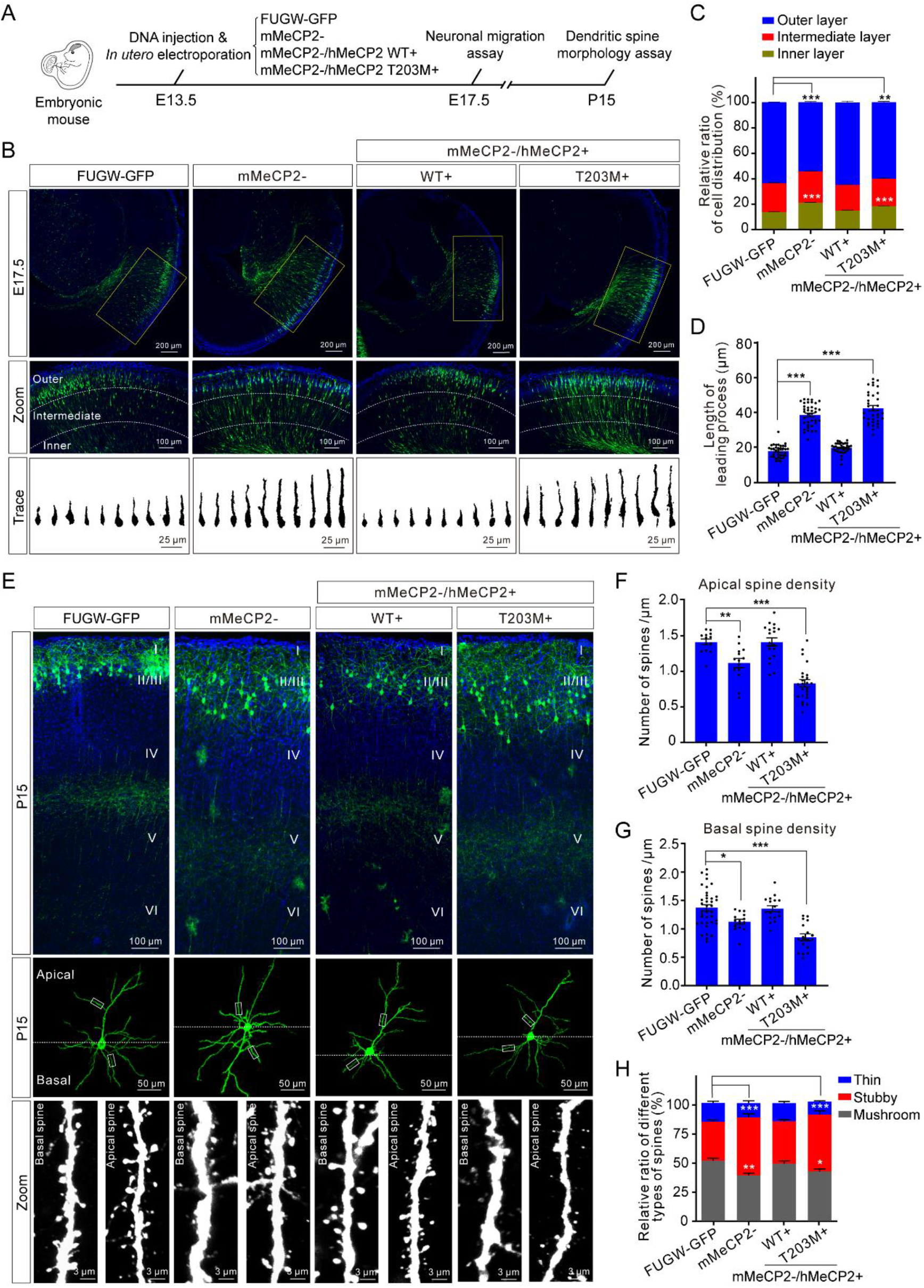
T203 O-GlcNAcylation is Required for Dendritic Spine Morphogenesis *in vivo*. (A) Schematic of the experimental design. The indicated plasmids were *in utero* electroporated in the neocortex of E13.5 mouse embryos. The electroporated region was then isolated at E17.5 or P15 for neuronal migration assay or dendritic spine morphogenesis assay, respectively. (B) Distribution of GFP+ pyramidal neurons in the indicated plasmid electroporated neocortex at E17.5. Boxed areas of GFP+ neurons are shown at higher magnification to illustrate the detailed distribution within the neocortex. The neocortex was equally divided into three parts from the inside out showing as the inner layer, the intermediate layer, and the outer layer (Zoom panels). The representative single GFP+ neuron from indicated groups was traced and illustrated by Fiji software (Trace panels). Scale bars represents 200 μm, 100 μm, and 25 μm, respectively. (C) Quantification of the relative ratio of GFP+ neuron distribution (%) in distinct neocortical layers. Histograms show mean ± SEM. χ^2^-test, **p < 0.01, ***p < 0.001. (D) Quantification of the length of leading process (LP) in GFP+ neurons electroporated with the indicated plasmid. Histograms show mean ± SEM. One-way ANOVA followed by Bonferroni test, ***p < 0.001. (E) Representative images of dendritic spines on apical or basal dendrites of GFP+ neurons at P15 after electroporation with the indicated plasmid. Boxed areas of basal and apical dendritic fragments are shown at higher magnification to illustrate the detailed dendritic spine morphology (Zoom panels). Scale bar represents 100 μm, 50 μm, and 3 μm, respectively. (F-G) Quantification of the dendritic spine density on apical and basal dendrites in GFP+ neurons, respectively. Histograms show mean ± SEM. One-way ANOVA followed by Bonferroni test, *p < 0.05, **p < 0.01, ***p < 0.001. (H) Quantification of the distribution of three subtypes of dendritic spines in GFP+ neurons at P15. Histograms show mean ± SEM. χ^2^-test, *p < 0.05, **p < 0.01, ***p < 0.001.

Previous studies has shown that the leading process (LP) branch is a critical determinant for nuclear translocation during neuronal migration ^49^. Therefore, we analyzed the morphology and projection of LP in MeCP2 knockdown (mMeCP2-) and rescue neurons (mMeCP2−/hMeCP2+). Interestingly, we found that the length of LP in confirmed mMeCP2- neurons is much longer compared with FUGW-GFP control neurons (38.94 ± 1.07 μm in mMeCP2- neurons and 18.18 ± 0.62 μm in FUGW-GFP control neurons, p < 0.001, n = 35 and 37, respectively).

Moreover, the abnormal LP branches in confirmed mMeCP2- neurons can be significantly rescued by exogenous expression of WT hMeCP2, but not hMeCP2 T203M mutant (19.98 ± 0.49 μm in mMeCP2−/hMeCP2+ neurons and 42.76 ± 1.53 μm in mMeCP2−/hMeCP2 T203M+ neurons, p = 0.4528 and p < 0.001, respectively, compared with FUGW-GFP control, n = 36 and 35, respectively) (Fig. 5D). Collectively, these results indicate that the abnormally developed LP branches may be responsible for the migration defects in mMeCP2- neurons, and that hMeCP2 T203 O-GlcNAcylation is critical for maintaining normal LP branch morphology and the navigation of migrating neuronal precursors *in vivo*.

Dendritic spines have been showed to be abnormal in the cerebral cortex in RTT patients and *Mecp2*-null mice ^11, 13, 47, 50^. Thus, we next asked whether MeCP2 T203 O-GlcNAcylation is essential for postnatal dendritic spine morphogenesis *in vivo*. First, we analyzed both dendrite and dendritic spine morphology in Layer II/III cortical projection neurons on P15 mice, which had been manipulated by *in utero* electroporation as described above on E13.5 (Fig. 5A). GFP fluorescence was used to trace the morphology of entire neurons, including the apical and basal dendritic arbors. Initial analysis did not find striking differences based on gross morphology of the dendrites including polarity and dendritic orientation in individual groups of electroporated neurons (Fig. 5E). Next, we imaged apical and basal dendritic segments at high magnification for each groups of electroporated neurons (Fig. 5E), and found that the density of both apical and basal spines were significantly decreased in mMeCP2- neurons compared with FUGW-GFP control neurons (1.118 ± 0.065 /μm and 1.13 ± 0.041 /μm in mMeCP2- neurons; 1.414 ± 0.042 /μm and 1.37 ± 0.054 /μm in FUGW-GFP control neurons, p = 0.0045 and 0.013, respectively, n = 13)(Fig. 5F and G). Meanwhile, the spine formation defects in mMeCP2- neurons can be significantly rescued by ectopic expression of WT hMeCP2, but not by hMeCP2 T203M mutant (1.415 ± 0.057 /μm and 1.36 ± 0.052 /μm in mMeCP2−/hMeCP2+ neurons, and 0.831 ± 0.052 /μm and 0.85 ± 0.059 /μm in mMeCP2−/hMeCP2 T203M+ neurons, p = 0.99 and p < 0.001, respectively, compared with FUGW-GFP control neurons, n = 17) (Fig. 5F and G).

Dendritic spines are morphologically heterogeneous within the neocortex, and these differences between spines reflect functional viability and pathological states ^51^. To interrogate the effect of hMeCP2 T203 O-GlcNacylation on spine morphology, we next classified spine shapes into thin, mushroom, and stubby categories as previously defined ^52^. We found significantly fewer mushroom spines in mMeCP2- neurons at P15, compared with FUGW-GFP control neurons (38.79 ± 1.89 % in mMeCP2- neurons and 51.18 ± 2.03 % in FUGW-GFP control neurons, p = 0.001, n = 17 and 38, respectively) (Fig. 5H). In contrast, no changes in spine morphology were observed in mMeCP2−/hMeCP2+ rescue neurons, compared with FUGW-GFP control neurons (48.76 ± 2.27 % in mMeCP2−/hMeCP2+ neurons and 51.18 ± 2.03 % in FUGW-GFP control neurons, p = 0.9993, n = 18 and 38, respectively). However, mMeCP2−/hMeCP2 T203M+ neurons had significantly fewer mushroom spines with relatively increased number of stubby spines (41.96 ± 2.31 % mushroom type and 48.50 ± 3.11 % stubby type in mMeCP2−/hMeCP2 T203M+ neurons; 51.18 ± 2.03 % mushroom type and 32.90 ± 1.58 % stubby type in FUGW-GFP control neurons, p = 0.0465 and p < 0.001, respectively, n = 18 and 38, respectively) (Fig. 5H). Therefore, our results indicate that hMeCP2 T203 O-GlcNAcylation plays a critical role in the regulation of dendritic spine morphogenesis. Notably, the features displayed in mMeCP2−/hMeCP2 T203M+ cortical neurons nicely mimic the defects in dendrite and spine maturation and synaptogenesis in RTT patients ^11, 13, 50, 53^.

### T203 O-GlcNAcylation Regulates Excitatory Synaptic Transmission

Previous studies have shown that shRNA-mediated MeCP2 knockdown affects homeostatic synaptic plasticity in hippocampal neurons ^54, 55^. Long-term potentiation (LTP) is also impaired at excitatory synapses of layers II/III and V in the primary somatosensory cortex of *Mecp2*-null mutant mice ^56, 57^. However, the bases of these deficits remain unclear. To investigate whether synaptic transmission is regulated by hMeCP2 T203 O-GlcNAcylation, we treated neurons in layers II/III of the developing cortex with various LEMPRA constructs using *in utero* electroporation, and recorded from GFP-positive neurons on P21 (Fig. 6A). First, we recorded miniature excitatory postsynaptic currents (mEPSCs) in FUGW-GFP control and mMeCP2-projection neurons (Fig. 6B and 6C). The mEPSC amplitudes remained similar between mMeCP2- neurons and FUGW-GFP control neurons (10.23 ± 0.57 pA in FUGW-GFP control, n = 10 neurons, and 11.24 ± 0.26 pA in mMeCP2-, n = 9 neurons, p = 0.7859). Exogenous expression of WT or T203M mutant hMeCP2 in mMeCP2- neurons has on significant effect on the mEPSC amplitudes (12.71 ± 0.77 pA in mMeCP2−/hMeCP2+, n = 10 neurons, and 12.59 ± 0.88 pA in mMeCP2−/hMeCP2 T203M+, n = 8 neurons, compared with FUGW-GFP control neurons, p = 0.0939 and 0.1793, respectively) (Fig. 6D and 6E). In contrast, in agreement with the decreased dendritic spine density in mMeCP2- neurons (Fig. 4E, 5F and 5G) and previous report ^47^, the mEPSC frequency significantly decreased from 3.00 ± 0.26 Hz in FUGW-GFP control neurons to 1.62 ± 0.11 Hz in mMeCP2- neurons (n = 10 and 9 neurons respectively, p = 0.0048) (Fig. 6F and 6G). In addition, the reduced frequency of mEPSCs in mMeCP2- neurons was efficiently rescued by exogenous expression of WT hMeCP2, but not the hMeCP2 T203M mutant (2.81 ± 0.26 Hz in mMeCP2−/hMeCP2+, n = 10 neurons, compared with FUGW-GFP control neurons, p > 0.9999; 1.66 ± 0.12 Hz in mMeCP2−/hMeCP2 T203M+, n = 8 neurons, compared with FUGW-GFP control and mMeCP2−/hMeCP2+ neurons, p = 0.0093 and 0.0259, respectively) (Fig. 6F and 6G), further indicating that T203 O-GlcNAcylation is involved in the regulation of postsynaptic spine density and excitatory synaptic transmission *in vivo*.

**Figure 6.**
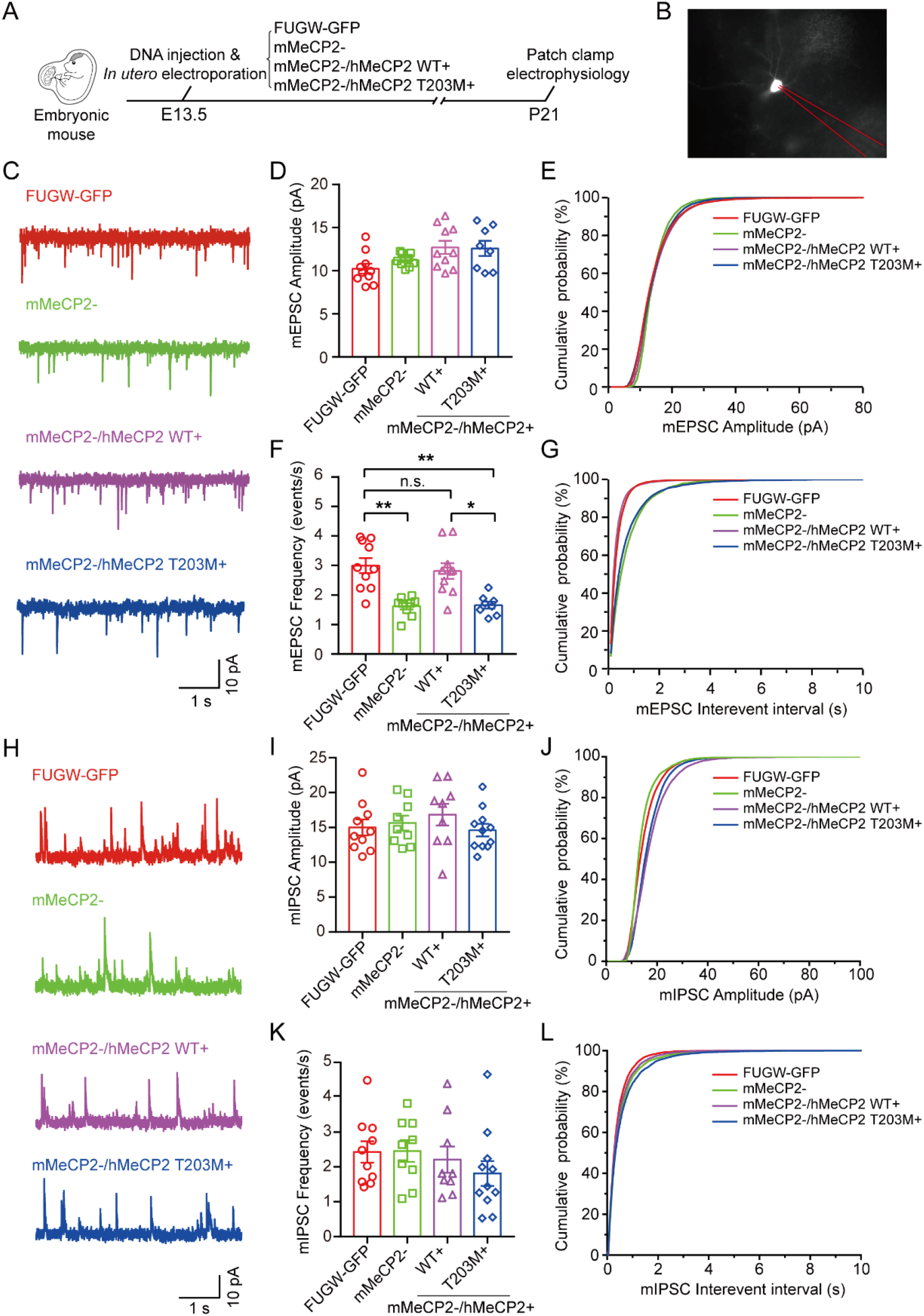
T203 O-GlcNAcylation is Essential for the Excitatory Synaptic Transmission in the Neocortex. (A) Schematic of the experimental design. The indicated plasmids were *in utero* electroporated in the neocortex of E13.5 mouse embryos. The electroporated region was then isolated at P21, and coronal brain slices were prepared for Patch clamp recording. (B) A representative GFP+ recording cell electroporated with indicated plasmid showing in A. (C) Representative traces of mEPSC recorded in Layer II/III GFP+ neurons electroporated with indicated plasmids. (D-E) Quantification of the amplitude and cumulative distributions of mEPSC in recorded GFP+ neurons electroporated with indicated plasmid. Histograms show mean ± SEM. Kruskal-Wallis ANOVA followed by Dunn’s post-hoc test, p > 0.05 for all comparisons. (F-G) Quantification of the frequency and cumulative distributions of mEPSC in recorded GFP+ neurons electroporated with indicated plasmid. Histograms show mean ± SEM. Kruskal-Wallis ANOVA followed by Dunn’s post-hoc test, *p < 0.05, **p < 0.01, n.s., not significant. (H) Representative traces of mIPSC recorded in Layer II/III GFP+ neurons electroporated with indicated plasmid. (I-J) Quantification of the amplitude and cumulative distributions of mIPSC in recorded GFP+ neurons electroporated with indicated plasmid. Histograms show mean ± SEM. Kruskal-Wallis ANOVA followed by Dunn’s post-hoc test. No significant differences were detected among each group, p > 0.9999 for all comparisons. (K-L) Quantification of the frequency and cumulative distributions of mIPSC in recorded GFP+ neurons electroporated with indicated plasmid. Histograms show mean ± SEM. Kruskal-Wallis ANOVA followed by Dunn’s post-hoc test. No significant differences were detected among each group, p > 0.9999 for all comparisons.

Interestingly, both the amplitude and the frequency of miniature inhibitory postsynaptic currents (mIPSCs) were not significantly different between mMeCP2- neurons and FUGW-GFP control neurons (amplitude: 15.63 ± 1.07 pA in mMeCP2-, n = 9 neurons, and 15.01 ± 1.15 pA in FUGW-GFP control, n = 10 neurons, p > 0.9999; frequency: 2.455 ± 0.31 Hz in mMeCP2-, n = 9 neurons, and 2.426 ± 0.31 Hz in FUGW-GFP control, n = 10 neurons, p > 0.9999) (Fig. 6H-L). Moreover, neither amplitude nor frequency of mIPSCs was changed in mMeCP2−/hMeCP2+ or mMeCP2−/hMeCP2 T203M+ neurons compared with FUGW-GFP control neurons (amplitude: 16.85 ± 1.56 pA in mMeCP2−/hMeCP2+, n = 9 neurons, p > 0.9999, and 14.61 ± 0.89 pA in mMeCP2−/hMeCP2 T203M+, n = 11 neurons, p > 0.9999; frequency: 2.209 ± 0.38 Hz in mMeCP2−/hMeCP2+, n = 9 neurons, p > 0.9999, and 1.811 ± 0.36 Hz in mMeCP2−/hMeCP2 T203M+, n = 11 neurons, p = 0.8159) (Fig. 6H-L). Together, these results indicate that T203M O-GlcNAcylation may be essential for the establishment of excitatory, but not inhibitory, synaptic transmission during neurodevelopment in the cortex.

### Neuronal Activity-Induced Bdnf Transcription is Dependent on T203 O-GlcNAcylation

MeCP2 regulates the expression of thousands of genes during neural development. One of the most important target genes is brain-derived neurotrophic factor (BDNF) ^21, 22, 24^. Numerous studies have shown that BDNF is critical for dendritic spine morphogenesis, synaptic maturation, and synaptic plasticity ^58, 59, 60, 61^. Moreover, activity-induced release of BDNF modulates spine morphology in conjunction with spontaneous neurotransmitter release ^59^. Therefore, we asked whether the impaired dendritic branching, spine morphogenesis and synaptic transmission observed in mMeCP2−/hMeCP2 T203M+ neurons were due to impaired *Bdnf* transcription. To evaluate the effect of T203 O-GlcNAcylation on neuronal activity-induced *Bdnf* transcription, primary mouse cortical neurons were dissected from E15.5 embryos and infected with various LEMPRA lentivirus constructions as described above (Fig. 4A) at DIV 3 for 4 days. We then added tetrodotoxin (TTX), a sodium channel selective blocker, into the culture medium at a final concentration of 1 μM to block the production of action potentials for 12 hr. On DIV 8, the cultured cortical neurons were treated with potassium chloride (KCl) at a final concentration of 55 mM for 5 hr to trigger synchronous membrane depolarization ^22, 62, 63^. The neurons were then harvested for RNA extraction and real time quantitative RT-PCR assay to investigate the transcription of MeCP2 target genes (Fig. 7A).

**Figure 7.**
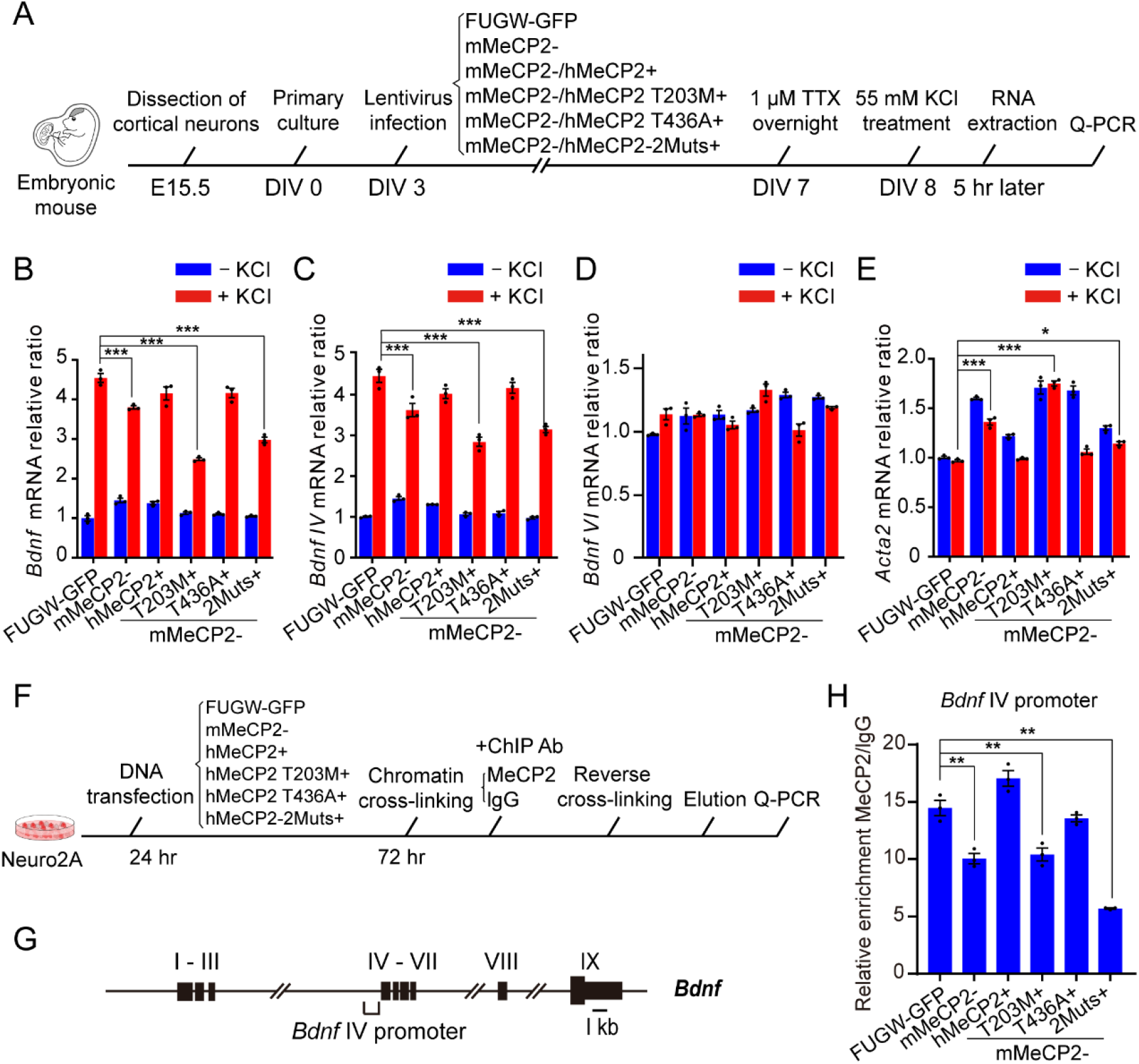
T203 O-GlcNAcylation Activates *Bdnf* Promoter IV-Dependent Transcription. (A) Schematic of the experimental design. The primary cortical neurons were isolated from E15.5 mouse embryos for *in vitro* culture. The neurons at DIV 3 were infected with the indicated lentivirus. The final concentration of 1 μM TTX was added in the culture medium at DIV 7. After 12 hr, the cultured cortical neurons were treated with 55 mM KCl for 5 hr to trigger synchronous membrane depolarization. Then the cultured neurons were harvested for RNA extraction and Q-PCR analysis. (B-E) Quantification of the transcription of total *Bdnf*, *Bdnf* exon IV, *Bdnf* exon VI, and *Acta2* at mRNA level in response to KCl treatment. Histograms show mean ± SEM. One-way ANOVA followed by Bonferroni test, *p < 0.05, ***p < 0.001. (F) Schematic of the experimental design. The cultured Neuro2A cells were transfected with the indicated plasmids for rescue experiments, and 48 hr later, the transfected cells were collected for ChIP assay. (G) A schematic depicting the total nine exons and exon IV promoter within *Bdnf* locus. (H) Quantification of the ChIP assay. The binding activity of *Bdnf* exon IV promoter to MeCP2 in indicated transfected cells were measured by Q-PCR amplification. Histograms show mean ± SEM. One-way ANOVA followed by Bonferroni test, **p < 0.01.

The transcription of total mouse *Bdnf* and its exon IV, but not exon VI, was dramatically enhanced by more than 4-fold in FUGW-GFP cortical neurons after KCl treatment (p < 0.001, n = 3) (Fig. 7B-D). The upregulation of total *Bdnf* and exon IV mRNA transcription were significantly lower in mMeCP2- neurons compared with FUGW-GFP control neurons after treatment with KCl (p < 0.001 and p < 0.001, respectively, n = 3) (Fig. 7B and C). Interestingly, the compromised upregulation of *Bdnf* transcription in mMeCP2- neurons could be significantly rescued by exogenous expression of WT hMeCP2 and hMeCP2 T436A mutant (p = 0.1015 and 0.1158, respectively, n = 3), but not with hMeCP2 T203M or hMeCP2-2Muts mutants (p < 0.001 and p < 0.001, respectively, n = 3) (Fig. 7B and C). These results suggest that T203 O-GlcNAcylation of hMeCP2 is essential for neuronal activity-induced enhanced transcription of *Bdnf*, especially for exon IV, but not exon VI. In addition, we examined the transcription of *Acta2*, a downstream gene normally repressed by MeCP2 in cortical neurons ^43^. We found that *Acta2* mRNA level was significantly upregulated in mMeCP2- neurons in both absence and presence of KCl treatment conditions (p < 0.001, n = 3) (Fig. 7E). Exogenous expression of WT hMeCP2 could significantly suppress the upregulated transcription of *Acta2* in mMeCP2- neurons, but hMeCP2 T203M and hMeCP2-2Muts failed to reverse the transcriptional activation effect in mMeCP2- neurons (p > 0.9999 and p < 0.001, respectively, n = 3) (Fig. 7E), indicating that T203 O-GlcNAcylation of hMeCP2 is also required to suppress *Acta2* transcription. These results suggest that hMeCP2 T203 O-GlcNAcylation is particularly important for activity-induced *Bdnf* expression in cortical neurons.

Finally, we performed chromatin immunoprecipitation (ChIP) analysis to quantify and compare the binding affinity of WT hMeCP2 and various hMeCP2 mutants to the mouse *Bdnf* IV promoter in mouse Neuro2A cells (Fig. 7F and G). As expected, binding of MeCP2 to the *Bdnf* IV promoter was significantly decreased in mMeCP2-cells (Fig. 7H). Exogenous expression of either WT hMeCP2 or hMeCP2 T436A mutant dramatically rescued the binding deficiency in mMeCP2-cells (p = 0.013 and 0.5981, respectively, compared with FUGW-GFP control cells, n = 3). In contrast, both hMeCP2 T203M and hMeCP2-2Muts failed to rescue the decreased binding of *Bdnf* promoter IV in mMeCP2-cells (p = 0.0004 and < 0.001, respectively, compared with FUGW-GFP control cells, n = 3) (Fig. 7H), suggesting decreased binding capacity of hMeCP2 T203M, but not T436A, to *Bdnf* promoter IV. Taken together, these results indicate that MeCP2 directly controls mouse *Bdnf* transcription by binding to the promoter IV. T203 O-GlcNAcylation on hMeCP2 is required to maintain *Bdnf* IV promoter binding, and therefore plays a critical role in the upregulation of *Bdnf* transcription following neural activation.

## DISCUSSION

Collectively, our results indicate that O-GlcNAcylation on hMeCP2 at T203 is critical for dendritic growth, spine formation, baseline and induced synaptic transmission, and the regulation of activity-induced *Bdnf* transcription. Using a LEMPRA-based shRNA system with exogenous expression of various hMeCP2 O-GlcNAc site mutants, we demonstrated that O-GlcNAcylation regulates MeCP2 activity during neurodevelopment, and pinpointed T203 as a major novel PTM site for O-GlcNAc modification on hMeCP2. Given that T203 mutations were previously implicated in clinical cases of RTT ^37, 38, 39^, these results suggest that hMeCP2 T203 O-GlcNAcylation not only supports normal cortical neurodevelopment, but its deregulation may contribute to the pathogenesis of RTT.

Mutations of the *MECP2* gene are the most prevalent cause of RTT, and some mutated sites are substrates for PTM ^38, 64, 65, 66^. This suggests that PTMs are a potentially crucial component of maintaining normal MeCP2 function, and deregulation may result in neurodevelopmental pathologies. Extensive studies have previously shown that phosphorylation ^21, 22, 23, 24^, acetylation ^67, 68^, SUMOylation ^64^ and other types of PTMs ^38^ can control MECP2 function, suggesting diverse forms of regulation.

Of particular interest was the finding that rat MeCP2 proteins are O-GlcNAcylated ^35, 36^, but its functional significance and mechanism was unclear. Previous results have implicated O-GlcNAcylation in the pathogenesis of neurodegenerative diseases ^69, 70^. For example, the imbalance of O-GlcNAcylation relative to phosphorylation on microtubule-associated protein Tau may be highly involved in the pathogenesis of Alzheimer’s disease ^71^. In addition, OGT-mediated O-GlcNAcylation modulates both the maturity and function of excitatory synapses in the developing brain ^72^, and changes in OGT levels may contribute to synaptic plasticity deficits during brain aging ^34^. Here, our results further illustrate a novel role for O-GlcNAcylation as a regulator of MeCP2 function during neurodevelopment. In particular, we identified T203 as a critical site for the establishment and maintenance of normal dendritic and spine development and synaptic transmission, in that exogenous expression of its mutant form T203M was not able to rescue deficits seen with knockdown of mouse endogenous MeCP2, whereas rescue effects were observed with other putative O-GlcNAcylation sites including S68, S204 and T436. The unique role of T203 may be due to its location on the human MeCP2 protein. MeCP2 can be subdivided into five domains corresponding to the N-terminal domain (NTD), the methyl-CpG binding domain (MBD), the intervening domain (ID), the transcriptional repression domain (TRD), and the C-terminal domain (CTD). By analyzing the *MECP2* mutation database, Bellini et al. have reported that about 25% of residues in the CTD and ID have been associated with pathogenic missense mutations in RTT ^38, 73^. In addition, the ID domain has been shown to be involved in MeCP2-mediated multiple protein-protein interactions as well as diverse phosphorylation events ^73^. Interestingly, T203 is located within the ID domain (Fig. 1B). Therefore, it is possible that the O-GlcNAcylation on T203 may affect its binding to DNA or its protein partners, resulting in subsequent regulation of neurodevelopment. However, the precise biological partners of T203 remain to be further identified. Our results showed that the T203M mutation had no effect on its binding to histone deacetylase 1 (HDAC1) (Fig. S5), but did result in reduced binding to the *Bdnf* IV promoter (Fig. 7). This suggests an HDAC1-independent transcriptional activation mechanism may underlie T203 O-GlcNAcylation in cortical neurons.

Although increasing evidence points to O-GlcNAcylation as a mediator of neurodegeneration and a modulator of neuronal signaling pathways in the brain ^27, 32, 71, 74, 75, 76, 77, 78, 79^, the underlying molecular mechanisms that support these crucial functions are not yet fully understood. Previously, the O-GlcNAcylation of CREB, a common upstream regulator of *Bdnf*, was found to critically regulate neuronal gene expression, axonal and dendritic growth, and long-term memory ^76, 79^. MeCP2 associates with CREB1 at the promoter of an activated target but not a repressed target ^80^. In addition, CREB signaling has also recently been shown to be involved in Rett syndrome pathogenesis ^81^. Interestingly, hMeCP2 T203 O-GlcNAcylation, but not its T203M mutant was able to rescue the activity-dependent *Bdnf* transcription (Fig. 7B). Because T203M dramatically reduces O-GlcNAcylation of hMeCP2, it suggests that O-GlcNAcyated T203 is necessary to support *Bdnf* transcription following neural depolarization. Interestingly, T203M did not notably impact *Creb* expression levels following KCl treatment (data not shown). This suggests that neuronal activation stimulates a variety of responses at both the O-GlcNAcylation PTM level and gene expression level to control neural development. First, both MeCP2 and CREB proteins are dynamically regulated via O-GlcNAcylation to stimulate normal dendrite and spine growth ^76^. Second, neural activity may result in the increased association of the MeCP2 and CREB1 ^80^, and the coupling between T203 O-GlcNAcylated MeCP2 with *Bdnf* exon IV, as well as the dissociation between phosphorylated MeCP2 with the *Bdnf* exon III promoter to promote *Bdnf* transcription ^21^. Together, our data and previous results indicate that upon neuronal activation, O-GlcNAcylation of MeCP2 upregulates the CREB-BDNF signaling pathway *via* multiple mechanisms to promote neural development and synaptic plasticity, with T203 as a direct binding partner of the *Bdnf* promoter.

Given the prevalence of PTMs on MeCP2, it is interesting to further consider the role of O-GlcNAc among other forms of PTM, especially in the context of RTT. Thus far, the best characterized PTM of MeCP2 protein is its phosphorylation ^21, 22, 23, 24^, and deregulation of MeCP2 phosphorylation may be involved in the pathogenesis of RTT ^65, 66^. Interestingly, O-GlcNAcylation has been closely linked to phosphorylation as both modifications can occur at the same or adjacent sites ^82^, and functional interaction between both modifications has also been previously characterized ^83^. However, in this study our MS assay did not identify the O-GlcNAc site T203 also as a phosphorylation site (Fig. 1 and Fig. S1), suggesting that the observed neurodevelopmental regulatory effects are likely due to O-GlcNAcylation on T203, rather than phosphorylation. In addition, we analyzed the effects of T203M deglycosylation on the phosphorylation of MeCP2 S421 and S80 ^22, 23^, but did not find any significant changes in phosphorylation level on either site (Fig. S6). Thus, it is possible that T203 O-GlcNAcylation affects neural development and activity-dependent transcription by independently recruiting a specific set of molecular coactivator to target gene promoters. Given the complexity of various genetic mutations in the pathogenesis of RTT ^38, 84^, and that some mutations impact PTM sites ^38^, delineating the effects of disease-related PTM sites and their interaction may lead to novel therapeutic targets for reversing neural deficiencies in RTT.

In this study, we identified several previously unknown O-GlcNAcylation sites in human MeCP2 and revealed novel functions of T203 O-GlcNAc modification in the regulation of neural development and synaptic transmission. Furthermore, we provided mechanistic insight into downstream signaling of MeCP2 T203 O-GlcNAcylation, adding to our understanding of the complex signaling network following MeCP2 PTMs in mediating neuronal activity-dependent transcription. Future work will need to further map out major relevant molecular pathways affected by MeCP2 T203 O-GlcNAcylation and their possible involvement in the pathogenesis of RTT disorders.

## Materials and Methods

### Animals

Pregnant ICR mice were purchased from SiBeiFu Co. (Beijing, China) and housed in the animal breeding facility of the Beijing Institute of Basic Medical Sciences. All animals were maintained and utilized in accordance with the guidelines of the Institutional Animal Care and Use Committee of the Beijing Institute of Basic Medical Sciences. *MECP2* Tg mice (JAX lab, #008679) ^19^ were kindly provided by Dr. Zilong Qiu at the Institute of Neuroscience, Chinese Academy of Sciences. *MECP2* Tg mice genotypes were determined by PCR assays using murine tail DNA, with primers as follows: 5’-CGCTCCGCCCTATCTCTGA-3’ (forward) and 5’-ACAGATCGGATAGAAGACTC-3’ (reverse).

### Plasmids

The rat *Mecp2* gene expression plasmid was a gift from Dr. Zilong Qiu at the Institution of Neuroscience, Chinese Academy of Sciences. The human *MECP2-e1* gene expression plasmid was a gift from Dr. Keping Hu at Chinese Academy of Medical Sciences & Peking Union Medical College. N-terminal HA-tagged *MECP2* and GFP-tagged *MECP2* CDS were subcloned into the pXJ40-HA and pEGFP-C1 vector, respectively. Other constructs of various *MECP2* mutations and indicated truncations were all generated based on the recombinant pXJ40-HA-hMeCP2 or pEGFP-hMeCP2 constructs. N-terminal His-tagged *MECP2* and GST-tagged *MECP2* CDS were subcloned into the pET-28a and pGEX-6P-1 vector, respectively. GST-tagged full length OGT was subcloned into the pET-28a plasmid. GST-OGT (323-1041) and GST-OGA (31-624) plasmids were gifts from Dr. Huadong Pei at National Center for Protein Sciences (Beijing). The PCR primers used for subcloning of truncated or mutant MeCP2 and OGT were listed in supplemental Table S3.

The MeCP2 LEMPRA (lentivirus-mediated protein-replacement assay) plasmid (named *pLenti-FUGW-shMeCP2-GFP-IRES-Flag-MECP2*) is a lentiviral vector with dual promoters. It was constructed by inserting the H1 promoter-driven mouse *Mecp2*-specific shRNA cassette against the sequence of 5’GTCAGAAGACCAGGATCTC-3’ into the indicated site ^85^, and inserting Flag-tagged shRNA-resistant human *MECP2* coding sequence under the control of the Ubiquitin-C (Ubc) promoter. The shRNA-resistant Flag-*MECP2* was generated by introducing five silent nucleotide mutations indicated in the following lower case letters within the coding sequence of *MECP2*: 5’-GagcGAAGACCAaGAcCTC-3’ ^24^.

### Cell culture and DNA transfection

HEK293T and Neuro2A cells were cultured in DMEM basic (Gibco, C11995500BT) and MEM basic (Gibco, C11095500BT) medium, respectively, supplemented with 10% FBS (Gibco, 10099-141C), 100 U/mL penicillin-streptomycin (Gibco, 15140122), in a 37°C incubator with a humidified, 5% CO_2_ atmosphere. Lipofectamine 2000 (Invitrogen, 11668019) was used for transfection following the manufacturer’s protocol.

### Mass spectrometry

HA-MeCP2 was ectopically expressed in HEK293T cells. Proteins were isolated by co-IP, and eluted using 200 μg/ml of HA peptide. The endogenous mouse MeCP2 was enriched by co-IP with anti-MeCP2 antibody (Cell Signaling Technology, 3456) and subjected to Sodium dodecyl sulfate-polyacrylamide gel electrophoresis (SDS-PAGE). The gel was stained with Coomassie brilliant blue. Visualized bands were excised, de-stained with ammonium bicarbonate buffer, and dehydrated in 75% acetonitrile. Following rehydration (with 50 mM ammonium bicarbonate), the gel slices were crushed and subjected to overnight digestion with trypsin or chymotrypsin. The peptides were extracted with acetonitrile containing 0.1% formic acid and vacuum dried. Proteolytic peptides were reconstituted with mobile phase A (2% acetonitrile containing 0.1% formic acid) and then separated on an on-line C18 column (75 μm inner diameter, 360 μm outer diameter, 10 cm, 3 μm C18). Mobile phase A consisted of 0.1% formic acid in 2% acetonitrile and mobile phase B was 0.1% formic acid in 84% acetonitrile. A linear gradient from 3 to 100% B over 75 minutes at a flow rate of 350 nL/min was applied. Mass spectrometry analysis was carried out on a Q-Exactive mass spectrometer (Thermo Fisher, SJ) operated in data dependent scan mode. Survey scan (m/z 375–1300) was performed at a resolution of 60,000 followed by MS2 scans to fragment the 50 most abundant precursors with collision induced dissociation. The activation time was set at 30 ms, the isolation width was 1.5 amu, the normalized activation energy was 35%, and the activation q was 0.25.

Mass spectrometry raw files were scanned and analyzed with the Proteome Discoverer software (version 2.1, Thermo Fisher Scientific) using MASCOT search engine with percolator against the human or rodent ref-sequence protein database. The mass tolerance was set to 20 ppm for precursor and 0.5 Da for product ion. Missed cleavages were no more than two for each peptide. O-GlcNAc of Ser/Thr were used as variable modifications.

### In vitro O-GlcNAcylation assay by chemoenzymatic labelling

Chemoenzymatic labelling and biotinylation of proteins in cell lysates were carried out following the manufacturer’s instructions. Briefly, WT and *MECP2* Tg mice brain lysates (200 μg) were labelled using the Click-iT O-GlcNAc Enzymatic Labelling System (Molecular Probes, C33368) protocol, then conjugated with an alkyne-biotin compound according to the Click-iT Protein Analysis Detection Kit (Molecular Probes, C33372) protocol. A parallel negative control experiment was performed in the absence of the labelling enzyme GalT or UDP-GalNAz. The methanol and chloroform were used to precipitate the biotinylated lysates. The biotinylated products were solubilized using 1% SDS solution, and neutralized with the neutralization buffer (6% Nonidet P40, 100 mM Na_2_HPO_4_, 150 mM NaCl, 50 mM Tris-HCl, pH 7.5, and protease inhibitor cocktail). Lysates were then incubated with streptavidin resin with end-to-end rotation at 4°C overnight. Resin was then washed five times with low-salt buffer (100 mM Na_2_HPO_4_, 150 mM NaCl, 0.1% SDS, 1% Triton X-100, 0.5% sodium deoxycholate) and five times with high-salt buffer (100 mM Na_2_HPO_4_, 500 mM NaCl, 0.2% Triton X-100). Biotinylated proteins were boiled with SDS loading buffer and then resolved with SDS-PAGE and subjected to Western blot analysis using anti-MeCP2 antibody (Cell Signaling Technology, 3456). To quantify the level of O-GlcNAcylation, the intensity of the total MeCP2 protein band (Input) and the O-GlcNAc MeCP2 protein band (Elution) were measured, and the ratio of the intensity of the O-GlcNAc protein versus the intensity of the total protein was taken as the level of O-GlcNAcylation ^86^.

### Primary culture of cortical neurons

Cortical neurons and hippocampal neurons were dissected and cultured from E15.5 and E17.5 embryonic mouse brain respectively, and neurons were maintained in Neurobasal medium (Gibco, 21103-049) supplemented with 2% B27 (Gibco, 17504044) and 1 mM GlutaMAX (Gibco, 35050061), and 100 U/mL penicillin-streptomycin (Gibco, 15140122). Cells were typically seeded at a density of 1-3 × 10^5^ cells/cm^2^ on dishes coated with poly-L-lysine (Sigma-Aldrich, P1524). Neuronal cultures were treated overnight in 1 μM tetrodotoxin (TTX) (Kangte Biotech, purity > 99%; 121206) to reduce endogenous neuronal activity prior to stimulation. Neurons were membrane depolarized with 55 mM extracellular KCl as previously described ^21^.

### Lentivirus package and purification

Lentiviruses were produced by co-transfection of HEK293T cells with the MeCP2 LEMPRA plasmid and the helper plasmids psPAX2 and VSV-G. Lentiviruses were concentrated by ultra-centrifugation 48-72 hr after transfection, and viral titers were determined by infection of HEK293T cells and determined by qPCR. Primary cultured neurons were infected with the purified lentivirus at appropriate time at 1 × 10^6^ TU/mL.

### Real time quantitative RT-PCR

Total RNA was extracted using TRIzol (Invitrogen, 15596018). 0.5 μg of RNA was used for reverse transcription using the Reverse Transcription System (TaKaRa, RR036) according to the manufacturer’s protocol. Real-time PCR was conducted in triplicates employing SYBR Green PCR master mix (CWBIO, CW2601) with the appropriate forward and reverse primers. For quantitative analysis of gene expression, results were averaged from three replicates in three independent experiments. Values were normalized to *Actb* levels. All PCR reactions were performed in triplicate with the following primers. *Bdnf* total: 5’-TGCCTAGATCAAATGGAGCTTCTC-3’ (Forward) and 5’-CCGATATGTACTCCTGTTCTTCAGC-3’ (Reverse); *Bdnf* exon IV: 5’-CAGAGCAGCTGCCTTGATGTT-3’ (Forward) and 5’-GCCTTGTCCGTGGACGTTTA-3’ (Reverse); *Bdnf* exon VI: 5’-GGGATCCGAGAGCTTTGTGTGGA-3’ (Forward) and 5’-GTAGGCCAAGTTGCCTTGTCCGT-3’ (Reverse); *Acta2*: 5’-GAGCTACGAACTGCCTGACG-3’ (Forward) and 5’-TACCCCCTGACAGGACGTTG-3’ (Reverse); *Actb*: 5’-GGCTGTATTCCCCTCCATCG-3’ (Forward) and 5’-CCAGTTGGTAACAATGCCATGT-3’ (Reverse).

### Expression and purification of recombinant proteins

Both His-tagged and GST-tagged proteins were expressed in *Escherichia coli* BL21. Bacteria were treated with 0.1 mM Isopropyl β-D-thiogalactoside (IPTG) (Thermo Fisher, AM9462) at 16°C for 16 hr to induce protein expression. To purify the His-tagged recombinant protein, the induced bacteria were harvested and suspended in 10 mM PBS (pH 7.4) containing 20 mM imidazole (Sigma-Aldrich, I2399) and 1 mM Phenylmethanesulfonyl fluoride (PMSF) (Sigma-Aldrich, 10837091001), followed by ultrasonication. The recombinant protein in the supernatant was incubated with the Ni Magnetic beads for 2 hr at 4°C, washed in 10 mM PBS (pH 7.4) three times, eluted with 10 mM PBS (pH 7.4) containing 200 mM iminazole, and followed by dialysis with 10 mM PBS (pH 7.4). Quantification of protein amount was measured by Coomassie brilliant blue staining.

To purify GST-tagged recombinant proteins, induced bacteria were harvested and suspended in 10 mM PBS (pH 7.4) containing 1 mM PMSF, followed by ultrasonication. Recombinant GST-tagged protein in the supernatant was purified using glutathione-Sepharose 4B beads (GE Healthcare, 17-0756-01), washed in 10 mM PBS (pH 7.4) three times, eluted with 10 mM PBS (pH 7.4) containing 20 mM reduced glutathione, and followed by dialysis with 10 mM PBS (pH 7.4). Quantification of the protein amount was measured by Coomassie brilliant blue staining.

### In vitro O-GlcNAcylation assay

Purified recombinant GST-OGT fusion protein (323-1041) was incubated with wild type His-tagged recombinant hMeCP2 or various His-tagged hMeCP2 mutants in 50 μL reactions (50 mM Tris-HCl, 12.5 mM MgCl_2_, 2 mM UDP-GlcNAc 1 mM DTT, pH 7.5) for overnight at 37°C, and Western blot analysis was carried out with anti-O-GlcNAc (RL2) antibody (Abcam, ab2739). For O-GlcNAc cleavage assay, O-GlcNAcylated proteins were treated with purified recombinant GST-OGA (31-624) fusion protein for 2 hr at 37°C in a volume of 50 μL, and Western blot analysis was carried out with anti-O-GlcNAc (RL2) antibody (Abcam, ab2739). Quantitative analysis of recombinant protein was measured by Coomassie brilliant blue staining ^42^.

### GST Pull-down assay

Bacteria-expressed GST, GST-hMeCP2 or GST-OGT fusion proteins were immobilized on glutathione Sepharose 4B beads (GE Healthcare, 17-0756-01) and washed three times with GST binding buffer (50 mM Tris-HCl, pH 7.5, 100 mM NaCl, 50 mM NaF, 2 mM EDTA, 1% Nonidet P40, and protease inhibitor cocktail). The beads were next incubated with His-hMeCP2 or His-OGT recombinant protein lysates at 4°C for 4 hr under rotation. Beads were washed with GST binding buffer (50 mM Tris-HCl, pH 7.5, 100 mM NaCl, 50 mM NaF, 2 mM EDTA, 1% Nonidet P40, and protease inhibitor cocktail) and proteins were eluted, followed by Western blot with indicated antibodies.

### Co-immunoprecipitation (Co-IP) assay

For endogenous co-IP assay, mouse brain lysates or cell lysates (1-2 mg) with protease inhibitor cocktail were incubated with anti-MeCP2 (Cell Signaling Technology, 3456) or anti-OGT (Sigma-Aldrich, HPA030751) antibody overnight at 4°C. After incubation, protein-A/G agarose beads were used for precipitation for 2 hr. The precipitates were then washed 5 times with the lysis buffer and eluted by boiling in SDS sample buffer for Western blotting. The gel was transferred to PVDF membrane, and the membrane was blocked with 5% milk in TBST buffer for 1 hr (at room temperature). It was then incubated overnight at 4°C with antibody, washed three times in TBST, and the signals were revealed by HRP reaction using the SuperSignal Chemilluminescent Substrate (Beyotime, P0018AS).

For co-IP in HEK293T cells, HEK293T cells were plated on 60 mm plates, and transfection was performed when the cells reached 50% confluence. Lipofectamine 2000 was used for transfection. A total of 5 μg of DNA was used in 60 mm plates at a molar ratio of 1:1 for GFP-tagged and HA-tagged constructs. Cells were harvested 36 hr later. The cells were rinsed with cold PBS, harvested, and lysed for 20 min at 4°C in a modified RIPA lysis buffer. 10% of the supernatant was saved for the input control, and the rest was incubated with 2 μg anti-HA antibody (Cell Signaling Technology, 3724) or anti-GFP (Cell Signaling Technology, 2955) overnight at 4°C. The immune complex was isolated by addition of 30 ml of a 50% slurry of mixed protein-A/G agarose for 2 hr, washed three times with the lysis buffer, then eluted by boiling-SDS lysis, and resolved by 8% SDS-PAGE. The gel was transferred to PVDF membranes, and the membrane was blocked with 5% milk in TBST buffer for 1 hr at room temperature. It was then incubated overnight at 4°C with the antibody, washed three times in TBST, and the signals were revealed by HRP reaction using the SuperSignal Chemiluminescent Substrate (Beyotime, P0018AS).

### Chromatin immunoprecipitation (ChIP) assay

Chromatin immunoprecipitation (ChIP) was prepared using the Upstate Biotechnology and Abcam kit following the manufacturer’s protocol. Briefly, Neuro2A cells were cross-linking at room temperature by addition of 1% formaldehyde for 15 min. Cross-linking was stopped by addition of 0.2 M glycine for 5 min at room temperature. Cells were washed three times in 10 ml ice-cold 10 mM PBS (pH 7.4), re-suspended in lysis buffer (50 mM Tris-HCl at pH 8.0, 1% SDS, 5 mM EDTA, and protease inhibitors) and directly sheared by sonication and processed for ChIP assay. Lysates was precleared by incubating in 1 mL diluted chromatin with a salmon sperm DNA /protein A-Sepharose (50 μL 50% slurry in 10 mM Tris-HCl, pH 8.0, 1 mM EDTA) for 2 hr at 4 °C with agitation. Beads were pelleted using brief centrifugation and the supernatant fraction collected. The rest of the supernatant was divided in two fractions: one for an equivalent amount of normal rabbit IgG control and the second incubated with ChIP-grade anti-MeCP2 (Abcam, ab2828) at 4°C with agitation overnight. Salmon sperm DNA/protein A-Sepharose slurry was added to the immune complexes and incubated at 4°C for 2-4 hr. Sepharose beads were collected and washed sequentially for 10 min each in the following buffers: once in low salt buffer (0.1% SDS, 1% Triton X-100, 2 mM EDTA, 20 mM Tris-HCl pH 8.0, 150 mM NaCl), three times in high salt buffer (0.1% SDS, 1% Triton X-100, 2 mM EDTA, 20 mM Tris-HCl, pH 8.0, 500 mM NaCl), twice in LiCl buffer (250 mM LiCl, 1% Nonidet P40, 1% deoxycholate, 10 mM Tris-HCl pH 8.0, 1 mM EDTA), and twice in TE buffer (10 mM Tris-HCl pH 8.0, 1 mM EDTA, pH 8.0). Complexes were eluted using elution buffer (250 μl of 1% SDS and 100 mM NaHCO3), formaldehyde cross-links reversed and DNA was precipitated and re-suspended. Both input and immunoprecipitated samples were analyzed by quantitative RT-PCR with *Bdnf* promotor IV primers: 5’-GGTCTTTAAGGTGGCCCAAG-3’ (forward) and 5’-TGGAGCATGTGATCAAAACAA-3’ (reverse).

### In utero electroporation

*In utero* electroporation (IUE) of E13.5 pregnant mice (ICR; commercially obtained from SiBeiFu) was performed as previously described ^48^. Specifically, pregnant mice were anesthetized with isoflurane (Yuyanbio, Shanghai). The uterine horns were exposed by Cesarean section and sterile, pre-warmed saline was repeatedly applied during the operation to keep the intestines moist. Animals were kept on a heating pad during the entire operation. 1 μL of the DNA plasmids for electroporation (2 μg/μL) was injected into the lateral ventricle through a pulled glass capillary tube. DNA was electroporated into the neocortex. After electroporation, the uterine horns were carefully repositioned into the abdominal cavity, which was then filled with pre-warmed saline. Animals were left to recover in a clean cage and embryos allowed to continue their development. At E17.5 or P15, pregnant mice or young mice were sacrificed. The brains were dissected and post-fixed with 4% PFA overnight at 4°C. Coronal brain slices from IUE mice were prepared by Cryostat (Thermo Scientific, FSE) for immunofluorescent staining, or prepared by the automated vibrating blade microtome (Leica, VT1200 S) for the electrophysiological recording.

### Immunofluorescent staining

The immunofluorescent staining of frozen brain sections and cultured neurons was performed using standard techniques as previously described in our lab ^87, 88^. Briefly, frozen sections (30 μm) or cultured neurons were washed for 10 min with 0.5% Triton X-100/PBS (PBS-T) for three times and then blocked with 5% goat serum in PBST for 1 hr. The sections or cultured neurons were then incubated overnight at 4°C with primary antibodies, washed for 10 min with 0.5% PBS-T for three times and subsequently treated with Alexa Fluor 568- or Alexa Fluor 488-conjugated fluorescent secondary antibody (1:500; Biotium, 20103, 20012) 1 hr at room temperature. The nucleus was counterstained with DAPI (ZSGB-BIO, ZLI-9556) during mounting onto glass slides with anti-fade solution. All images were processed and analyzed using Olympus FV-1200 and Image J software.

### Electrophysiology

Preparation of brain slices was performed as previously described ^89, 90^. A single slice was then transferred to the recording chamber and submerged in a continuously flowing oxygenated NaHCO3-buffered saline (1.0-1.5 ml/min) warmed at 32°C. The recording electrodes had resistance of 3-5 MΩ when filled with internal solution consisting of (in mM): 135 mM cesium methanesulfonate, 10 mM Hepes, 0.2 mM EGTA, 8 mM NaCl, 4 mM Mg-ATP, and 0.3 mM Na3GTP (pH 7.2 with CsOH, osmolality adjusted to 280-290 mOsm). The slice was visualized with a 40 × water-immersion objective (Olympus) using standard infrared and differential interference contrast (IR-DIC) microscopy, and a CCD camera (QImaging, Surrey). Cells in the cortex up to ~60 μm beneath the slice surface were patched and monitored. Recording in normal voltage-clamp mode was performed with an Axon 700B amplifier (Molecular devices) and Clampex 10.5 software (Molecular devices). After tight-seal (>1 GΩ) formation, fast and slow capacitance compensation was auto performed. Neurons were excluded from the analysis when their series resistance was above 25 MΩ and changed by more than 25% during the experiment. Data were filtered at 2 kHz and acquired at a sampling rate of 10 kHz. For mEPSC recording, the target neurons were held at −70 mV in the presence of SR95531 (25 μM; Sigma-Aldrich, S106) and TTX (1 μM; Kangte Biotech, purity > 99%; 121206). For mIPSC recording, the target neurons were held at 0 mV in the presence of CNQX (25 μM; Sigma-Aldrich, C239), DL-AP5 (50 μM; Abcam, ab120271), and TTX (1 μM, Kangte Biotech, purity > 99%; 121206). 5 min consecutive miniature events were collected and analyzed. All mEPSCs or mIPSCs above a threshold value (5 pA) were included in the data analysis and each event was verified visually. Experiments were carried out in a genotype-blinded manner. No statistical analysis was used to predetermine the sample sizes used for experiments; however, our sample sizes are similar to those reported previously.

All drugs were purchased from Sigma-Aldrich (St. Louis, MI, USA) unless otherwise noted. Drugs were dissolved as concentrated stocks and stored at −20°C. Working solutions with different drugs were prepared just before use. During experiments, drugs were applied in the flowing bath solutions. Total replacement of the medium in the recording chamber occurred within 1 min. Data analysis was performed with software including Clampfit (Version 10.5), MiniAnalysis program (Version 6.0.3) Prism (Version 8.0), and Origin (Version 9.0).

### Dendritic length and spine density analysis

The length of dendritic branches in primary cultured hippocampus neurons was determined as follows: EGFP-positive neurons were randomly selected from each condition, and the dendritic length of all protrusions was analyzed using Fiji software. At least three independent experiments were performed, and the number of neurons >50 per condition were analyzed. For spine density analysis, confocal Z stacks of neurons in hippocampal were acquired with a confocal microscope (Olympus, FV-1200) using an oil-immersion 60 × objective lens. Images were analyzed with Fiji software. Protrusions in direct contact with the dendrites were counted as spines, and the average spine density was calculated as the number of spines per μm dendritic length. At least 500 μm dendrites from seven or more neurons were analyzed for each groups (more than 3 mice per group). All quantifications were analyzed with One-way analysis of variance (ANOVA).

### Statistical analysis

Data are presented as mean ± SEM and were analyzed by two-tailed Student’s t-test, or one-way ANOVA followed by Bonferroni test. Kruskal-Wallis ANOVA followed by Dunn’s post-hoc test was performed to analyze the data of electrophysiological experiments. A χ^2^test was applied to analyze the distribution of cells in either layers of the neocortex, and the distribution of different types of dendritic spines. Unless otherwise indicated, in figures, *p < 0.05, **p < 0.01, and ***p < 0.001. Changes were considered significant if the p value was < 0.05.

## SUPPLEMENTAL INFORMATION

Supplemental Information includes three tables and six figures can be found with this article as a separate supplementary file.

## AUTHOR CONTRIBUTIONS

H.W. designed the experiments. X.D. performed the MS experiment to identify the O-GlcNAcylation sites on MeCP2. J.C., L.C., R.D. and Y.W. performed the lentivirus packaging and purification, biochemical and molecular biology experiments. J.C. and Q.Z. performed the *in utero* electroporation experiments. Z.Z. and J.C. performed the electrophysiology experiments. H.W., X.D., M.F., J.C. and Z.Z. analyzed the results. H.W. and J.C. wrote the manuscript.

## ACKNOWLEDGMENTS

We thank Dr. Zilong Qiu at Institute of Neuroscience, CAS, Dr. Huadong Pei at the National Center for Protein Sciences (Beijing), and Dr. Ke-Ping Hu at Chinese Academy of Medical Sciences & Peking Union Medical College for providing the valuable reagents for this work, and thank Dr. Dong Yang at Beijing Proteome Research Center for his great help with the phylogeny tree analysis of MeCP2 protein sequence. We thank all members of the Wu laboratory for discussion. This work was supported by the National Natural Science Foundation of China (Grants 31770929 and 31522029), and the Beijing Municipal Science and Technology Commission (Grants Z181100001518001 and Z161100000216154) to H.W.

## CONFLICT OF INTEREST

The authors declare that they have no conflicts of interest with the contents of this article.

## Notes

### Competing Interest Statement

The authors have declared no competing interest.

